# Glucocorticoid signaling mediates CD8^+^ T cell memory differentiation

**DOI:** 10.1101/2022.07.19.500581

**Authors:** Azeez Tehseen, Dhaneshwar Kumar, Roman Sarkar, Sudhakar Singh, Abhishek Dubey, Sharvan Sehrawat

**Affiliations:** Department of Biological Sciences, Indian Institute of Science Education and Research (Mohali); SAS Nagar, Knowledge City, Sector 81, Mohali, 140306, Punjab

## Abstract

We provide evidence on the T_RM_ potentiation by glucocorticoids in stimulated CD8^+^ T cells. Signaling via glucocorticoid receptor reciprocally regulated short-lived effector cells and memory precursor effector cells. Influenza A virus infected mice treated transiently with dexamethasone preferentially generated multipotent T_RM_ cells that efficiently responded to the subsequent infection. Compromised memory following abrogation of glucocorticoid signaling further confirmed their role in memory potentiation. Transcriptomic and biochemical analysis of dexamethasone treated cells revealed a metabolic switch to oxidative phosphorylation via an engagement of AMPK signaling due to reduced glucose uptake. These cells exhibited an accumulation of phosphorylated STAT3 and STAT5 to drive memory differentiation. Therefore, glucocorticoids mediate tissue homing memory T cell differentiation.

**One-Sentence Summary:** Glucocorticoid signaling in responding CD8^+^ T cells promotes memory differentiation

## Main Text

CD8^+^ T cells (Cytotoxic T lymphocytes, CTLs) are an integral part of the vertebrate host’s defense against intracellular pathogens. Approximately 5-10% of the CTLs recruited during the acute phase differentiate into memory cells that can be rapidly recalled upon reinfection (*1, 2*). Based upon their migratory pattern and localisation, broadly three types of memory cell subsets are recognised, namely, central memory (T_CM_), effector memory (T_EM_) and tissue resident memory (T_RM_); each with largely common but some specialised functions (*3–6*). The non-migratory T_RM_ cells reside in non-lymphoid tissues and serve to restrict the virus at the site of infection. Since promoting T_RM_ responses could help optimize the efficacy of vaccines against local infections, considerable efforts are being made not only to define their phenotype but also to devise strategies that can boost such responses (*7*).

The infection induced micro-environment serves both to promote and control inflammation and can critically influence the differentiation program of CTLs (*8*). Although the role of tissue damaging events in maintaining the local immunity was shown recently, the influence of the generated microenvironment on local immune surveillance remains less well understood (*9*). Glucocorticoids (GCs) constitute one such class of molecules that have context dependent pro or anti-inflammatory roles (*10–12*). We had previously reported a preferential survival of effectors as compared to their quiescent counterparts following GC exposure (*13*). Since infections have been shown to induce the production of glucocorticoids and the sensing receptors are expressed by the CTLs, we undertook the study to investigate the influence of GC signaling in the differentiation of antigen-specific CD8^+^ T cells (*14*).

3×10^5^ OT1 (CD45.2^+^) cells were transferred into sex matched congenic (CD45.1^+^) mice and the recipients were infected with 200 pfu of influenza A virus (WSN-SIINFEKL) intranasally followed by dexamethasone and mifepristone treatment as indicated (Figure 1A). Longitudinal analysis of the donor cell kinetics revealed a dramatic increase in the frequencies in the circulation of the dexamethasone treated animals, with such cells being 10 fold more than those in the control animals even at 90 days post infection (dpi) (Figure 1B). An analysis at 7dpi revealed an enhancement of memory precursors amongst the responding donor cells in the circulation, lungs as well as lymphoid tissues of the dexamethasone treated animals, while pharmacological antagonism via mifepristone enhanced short lived effectors (Figure 2A & B). Therefore, GC signaling reciprocally regulated the differentiation of memory and short lived effectors. Upon a heterologous challenge at 90dpi with MHV68-SIINFEKL, the memory cells were efficiently recalled and an increased frequency of the donor cells was observed in lungs as well as the lymphoid organs of the dexamethasone treated animals (Figure 1C & D). The recalled cells in the dexamethasone group were functionally superior to the control cells as measured by intracellular cytokine staining (Figure 1E-H). Accordingly, upto a 30 fold increase in the counts of IFNγ, TNFα and GzmB producers were observed in the dexamethasone treated group. Moreover, the cells also exhibited a migratory phenotype, with a larger frequency of donor cells being CXCR3^+^, CD103^+^ and CD62L^lo^ (Figure S1A & B). Given the phenotype of tissue migrating anti-viral CD8^+^ T cells generated in the presence of GCs, we directly investigated their tissue homing potential. Tetramer positive cells were sorted from the draining mediastinal LN of sham and dexamethasone treated mice following an intranasal infection with WSN-SIINFEKL. The cells were labelled with two different concentrations of CFSE (CFSE^lo^- Sham T_x_ cells and CFSE^hi^- Dexa T_x_ cells), admixed in a 1:1 ratio and adoptively transferred into mice that were in the acute phase (6dpi) of WSN- SIINFEKL infection (Figure S2A). Two hours post transfer, the frequency of CFSE^hi^ and CFSE^lo^ cells was measured in different organs. CFSE^hi^ cells were four-fold more than CFSE^lo^ cells in lung tissues while a complete reversal in the ratios was observed in spleen and peripheral blood of the recipient animals (Figure S2). CFSE^lo^ and CFSE^hi^ cells were equally distributed in the LNs (Figure S2B and C). These results clearly showed that the virus-specific cells expanded under the influence of a transient dexamethasone therapy preferentially migrated to the infected tissue sites and establish a better memory. To test this, we analysed the distribution of donor cells in different organs at 210 dpi (Figure S3A). The donor cells were 10 times more in the different compartments (Figure S3C). The donor cells in the dexamethasone treated group exhibited better functionality (Figure S3D-F). Additionally, a higher proportion of the donor cells present in lungs, but not in the lymphoid tissues of the dexamethasone treated group stained for Ki67 (Figure S3G & H), indicative of their enhanced turnover at the non-lymphoid tissue (NLT). Therefore, not only did the dexamethasone therapy increase the basal level of long-term memory, it also endowed these memory cells with an better recallability and functionality. Therefore, GC signaling during the acute phase might contribute to the differentiation of tissue homing memory T cells.

**Figure 1.**
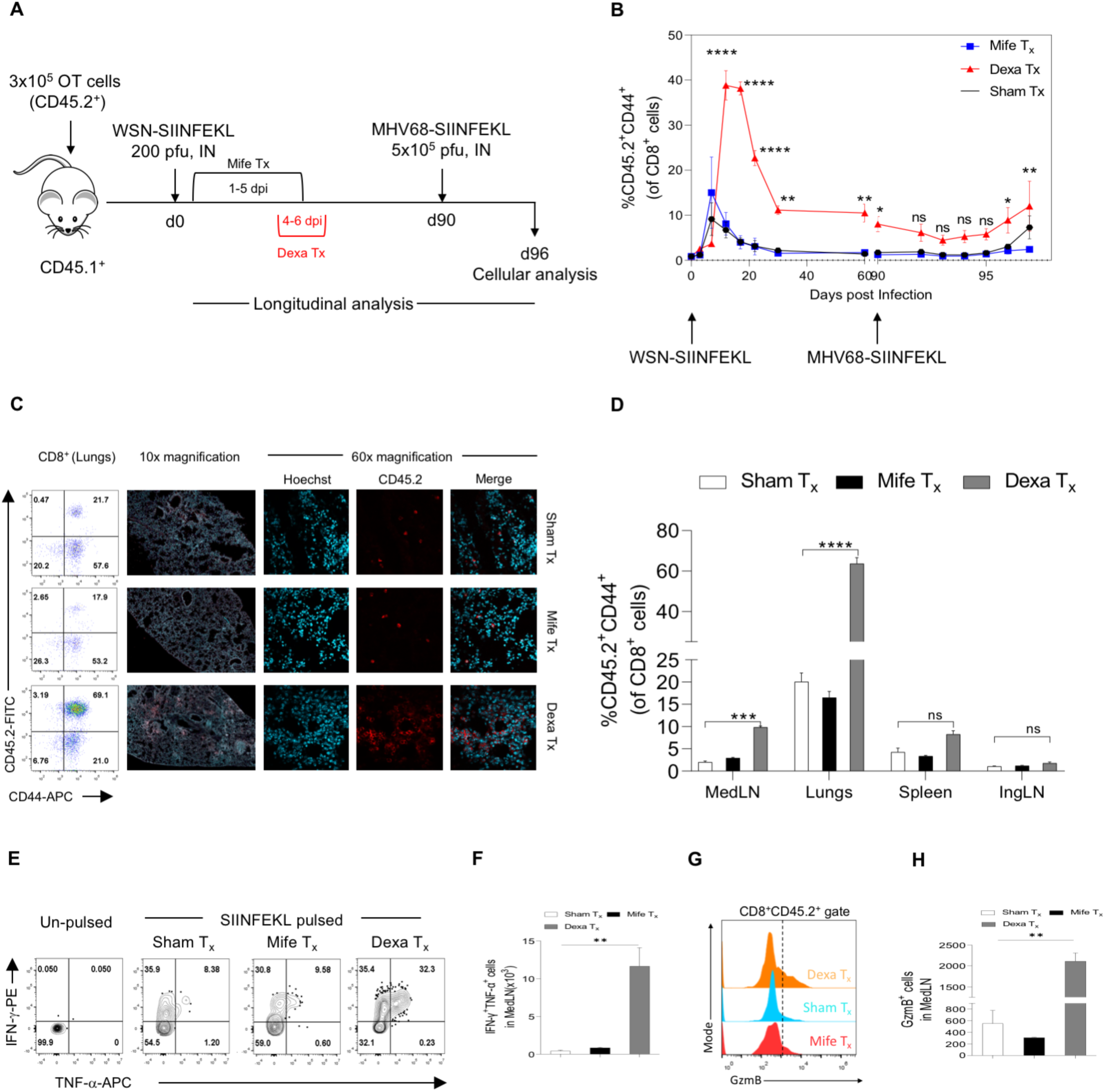
Dexamethasone mediated virus-specific CD8^+^ T cell memory enhancement. **A.** A schematic of the experiments is shown. 3×10^5^ OT-1 cells were transferred into naïve sex matched congenic CD45.1^+^ B6 mice. The recipients were then intranasally infected with 200 pfu of WSN-SIINFEKL. Randomized groups were intraperitoneally administered with either the diluent (Sham T_x_) or 40 mg/kg Body Weight (B Wt.) of mifepristone from 1-5 days post infection (dpi) or 10 mg/kg B. Wt. of dexamethasone from 4-6 dpi. The frequencies of the OT1 (CD45.2^+^) cells were then tracked in circulation at different days post infection (dpi). At 90 dpi the animals were intranasally infected with 10^5^ pfu of MHV68-SIINFEKL. The expansion of the donor (OT1) cells was also measured subsequently. **B.** The kinetics of expansion and contraction of the OT1 cells in the three groups are summarized. Data represents Mean ± SEM; ***p<0.001; **p<0.005; *p<0.05 and ns (p>0.05)- not significant (two-way ANOVA). Mice in each group were sacrificed at 7 days post recall and subjected to cellular analyses. **C**. Representative FACS plots and the confocal images from respective lung sections of each group are shown. **D**. Bar-diagram summarizing the frequency of donor cells in the various organs is shown. **E-H**. ICCS assays were performed to measure the functionality of the donor cells in each group after heterologous challenge. **E.** Representative FACS plots show the frequencies of donor cells producing IFN-γ and TNF-α. **F.** Bar diagrams summarizing the count of double positive donor cells producing IFN-γ and TNF-α is shown. **G.** Representative overlaid histograms for GzmB production by donor cells in the medLN of each group are shown. **H**. Bar diagram summarizing the count of GzmB producers in the medLN of each group is shown. The experiments were performed four times with similar results with n=3 per group. Data represents Mean ± SEM; ***p<0.001; **p<0.005; *p<0.05 and ns (p>0.05)- not significant (one-way ANOVA).

**Figure 2.**
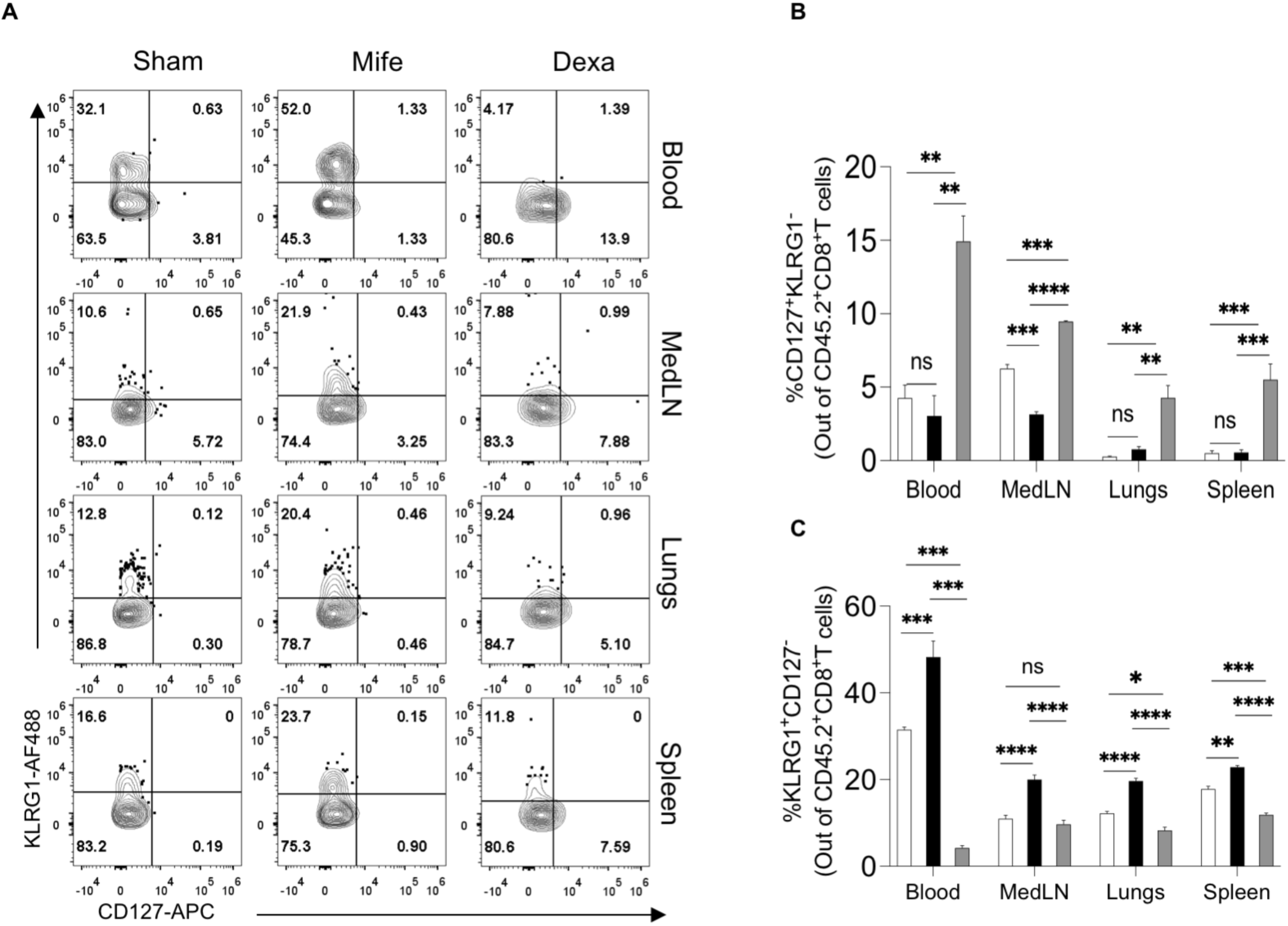
Dexamethasone enhances memory precursor phenotype. 10,000 OT-1 cells were transferred into naïve sex matched congenic CD45.1^+^ B6 mice. The recipients were then intranasally infected with 200 pfu of WSN-SIINFEKL. The animals were then treated with dexamethasone and mifepristone as indicated in Figure 1A. Cellular analysis was performed in the blood and various lymphoid and non-lymphoid organs at 7 dpi. **A.** Representative FACS plots for the analysis of MPECs (KLRG1^-^CD127^+^) and SLECs (KLRG1^+^CD127^-^) in different compartments are shown. **B & C.** Cumulative bar diagrams show the frequency of MPECs (**B**) and SLECs (**C**). The experiments were repeated three times with similar results with n=3 per group. Data represents Mean ± SEM; ***p<0.001; **p<0.005; *p<0.05 and ns (p>0.05)- not significant (Student’s T-test).

To show the role of T cell intrinsic programming in the GC mediated enhancement of T cell memory, we sorted the virus specific cells from WSN-SIINFEKL infected dexamethasone, mifepristone and sham treated animals at 7dpi and transferred these cells into naïve, unmanipulated mice (Figure 3A). A tracking of these cells in the circulation of the recipients revealed a slight increase in frequency of the tetramer positive cells in the circulation of Dexa T_x_ cell recipients 2 weeks after transfer. Upon heterologous challenge with MHV68-SIINFEKL virus, a rapid increase in virus-specific cells was observed in the circulation of the Dexa T_x_ cell recipients, with the tetramer positive cells ∼30 fold more in the circulation at 3 days post rechallenge (Figure 3B). The Dexa T_x_ cells were also ∼10 fold more frequent in the lung tissues, while these cells were barely detectable at other sites (Figure 3C and D). Additionally, the cells collected from lungs tissues of Dexa T_x_ recipients divided more efficiently following an *in vitro* peptide pulse as analysed by a CFSE dilution assay (Figure 3E & F). These results suggested for a dexamethasone induced programming of CD8^+^ T cells during their initial antigenic exposure that led to efficient recall, preferential homing and residence in non-lymphoid organs such as lungs. Both IFN-γ as well as IFN-γ and TNF-α producers were three fold more abundant in the animals that had received Dexa T_x_ cells in comparison to the control cells (Figure 3G and H). A significantly higher frequency of granzyme B producing Dexa T_x_ cells was also observed in the lungs as compared to the control cells (Figure 3I and J). Similar results were obtained when MHV68-SIINFEKL infection expanded CD45.2^+^OT1 cells exposed to dexamethasone, mifepristone and sham treatment were tracked following adoptive transfer in CD45.1^+^ congenic mice by homologous infection four months later (Figure S4). The recalled Dexa T_x_ cells controlled the replicating virus better as recipients of control cells had viral titres in excess of 100 fold in the lung tissues (Figure S4D).

**Figure 3.**
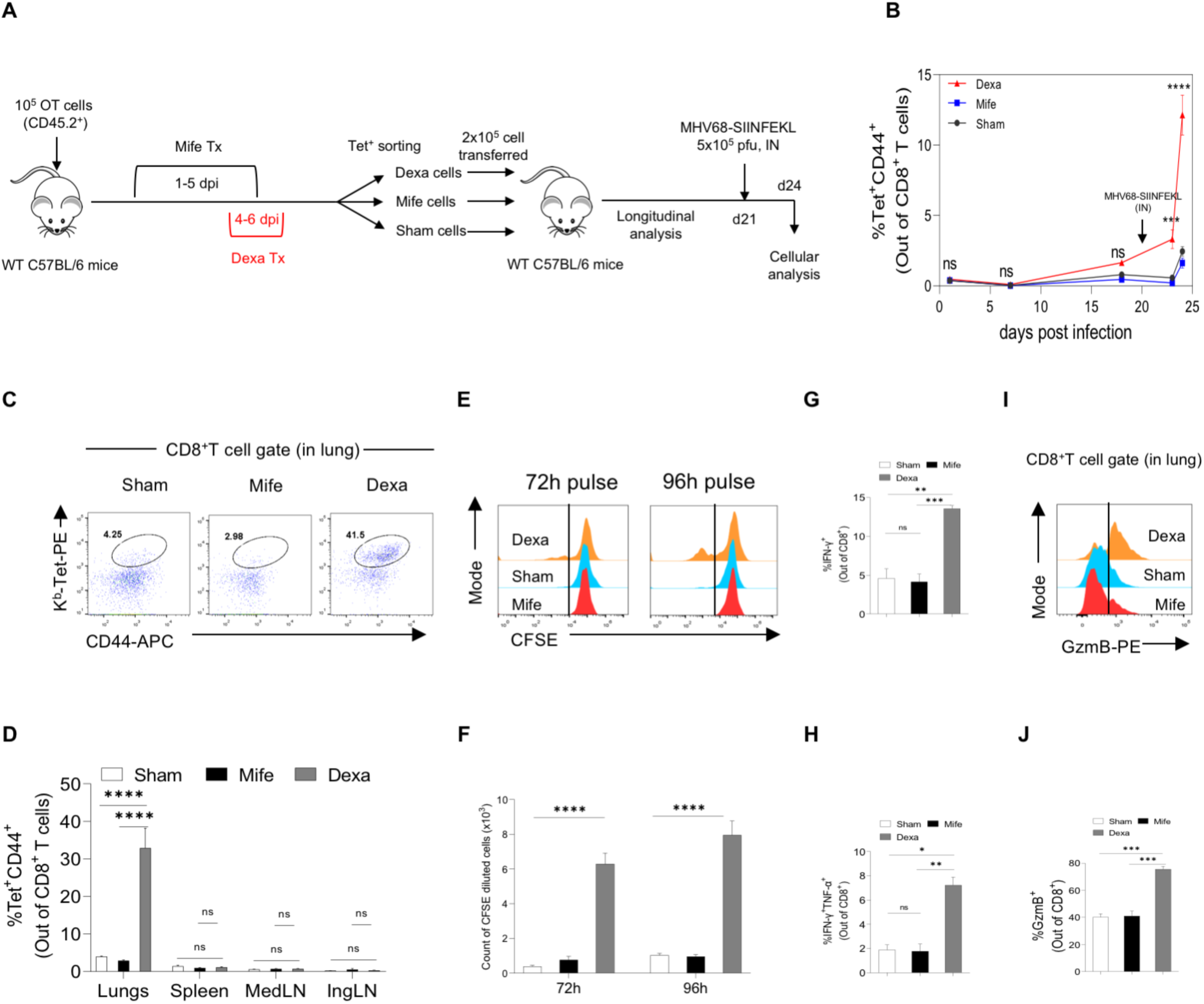
Enhanced survival and recall response of GC-exposed virus-specific CD8^+^ T cells. 10^5^ OT1 cells were transferred into naïve B6 mice which were then intraperitoneally infected with 200 pfu of WSN-SIINFEKL one day after the transfer and given the indicated treatment. 2.5×10^5^ SIINFEKL-Tetramer^+ve^ sorted from each of these groups at 7dpi were then transferred into naïve, sex-matched B6 mice. The fate of the transferred cells in each animal was then tracked at different days post transfer. 3 weeks post transfer, the animals were intranasally infected with 10^5^ pfu of MHV68-SIINFEKL. The kinetics of tetramer^+ve^ cells in circulation was then analyzed. The plot summarizing the kinetics of the transferred cells in circulation of recipients is shown. Data represents Mean ± SEM; ***p<0.001; **p<0.005; *p<0.05 and ns (p>0.05)- not significant (Two- way ANOVA). 3 days post reinfection, the mice were sacrificed and subjected to cellular analysis. **C**. Representative FACS plots depict virus-specific cells in the disrupted lung tissues. **D**. Bar diagrams summarize the frequency of K^b^-SIINFEKL-tetramer^+ve^ cells in the various organs analyzed**. E, F**. The proliferative potential of the virus-specific cells recovered from the lung tissues was analyzed by CFSE dilution assays. **E**. Representative half-offset histogram plots show the extent of CFSE dilution at 72 hours (left) and 96 hours (right) post peptide pulse. **F**. Bar diagrams summarize the count of CFSE diluted cells in each group at the mentioned time-points. Data represents Mean ± SEM; ***p<0.001; **p<0.005; *p<0.05 and ns (p>0.05)- not significant (Two-way ANOVA). **G, H**. Cytokine production by CD8^+^ T cells recovered from lung tissues in each group was measured by ICCS assay. Bar diagrams summarize the frequency of CD8^+^ T cells producing IFN-γ (**G**) and both IFN-γ and TNF-α (**H**) upon peptide stimulation. **I**. The representative overlaid histograms show the frequencies of GzmB producing CD8^+^ T cells recovered from lungs of animals in each group. **J**. Bar diagrams show the frequencies of GzmB producing CD8^+^ T cells. Data represents Mean ± SEM; ***p<0.001; **p<0.005; *p<0.05 and ns (p>0.05)- not significant (One-way ANOVA). The experiment was repeated twice with similar results with n=3 per group.

Finally, in order to unambiguously show the role of GR signalling in CD8^+^ T cell memory generation, we ablated the GR by shRNA mediated knockdown and analysed their response *in vivo*. We compared the phenotype of *nr3c1* depleted and control cells. The specific disruption of *nr3c1* by shRNA and not by the scrambled sequences was shown in the lysates of cells using anti- NR3C1 antibody (Figure S5A). A genetic deficiency of *nr3c1* downregulated the mRNA expression levels of CD127, CD103, IFN-γ and T-bet indicating their compromised memory differentiation and functionality (Figure S5B). *In vitro* stimulated control or the shRNA transduced OT1 cells were then tracked for their fate by transferring *nr3c1* depleted or the control OT1 cells into naïve C57BL/6 mice. The recipient animals were infected with WSN-SIINFEKL 14 days later (Figure 4). We observed reduced frequencies of expanded OT1 cells in the animals that had received *nr3c1* depleted cells (Figure 4A and B). At the peak of response, the frequency of K^b-^SIINFEKL-tetramer^+ve^ cells were three-fold lower in peripheral blood, draining medLNs and lung tissues of animals receiving knockdown cells as compared to those receiving control cells (Figure 4B). Additionally, a lower proportion of such cells in comparison to the control cells expressed CD69 and CD127 indicating for their inefficient activation and differentiation in the absence of GC signalling (Figure S5C-F). Consequently, the IFN-γ producers were less frequent among the knockdown cells as compared to the control cells following *ex vivo* stimulation with the cognate peptide (Figure 4C & D). Therefore, a transient exposure to dexamethasone during the acute phase of a primary infection resulted in enhanced memory transition as well as preferential localisation at the site of infection. Such cells conferred to the animals a quick protection following re-infection. Additionally, the responding antigen-specific CD8^+^ T cells depleted of the GR were compromised in memory differentiation.

**Figure 4.**
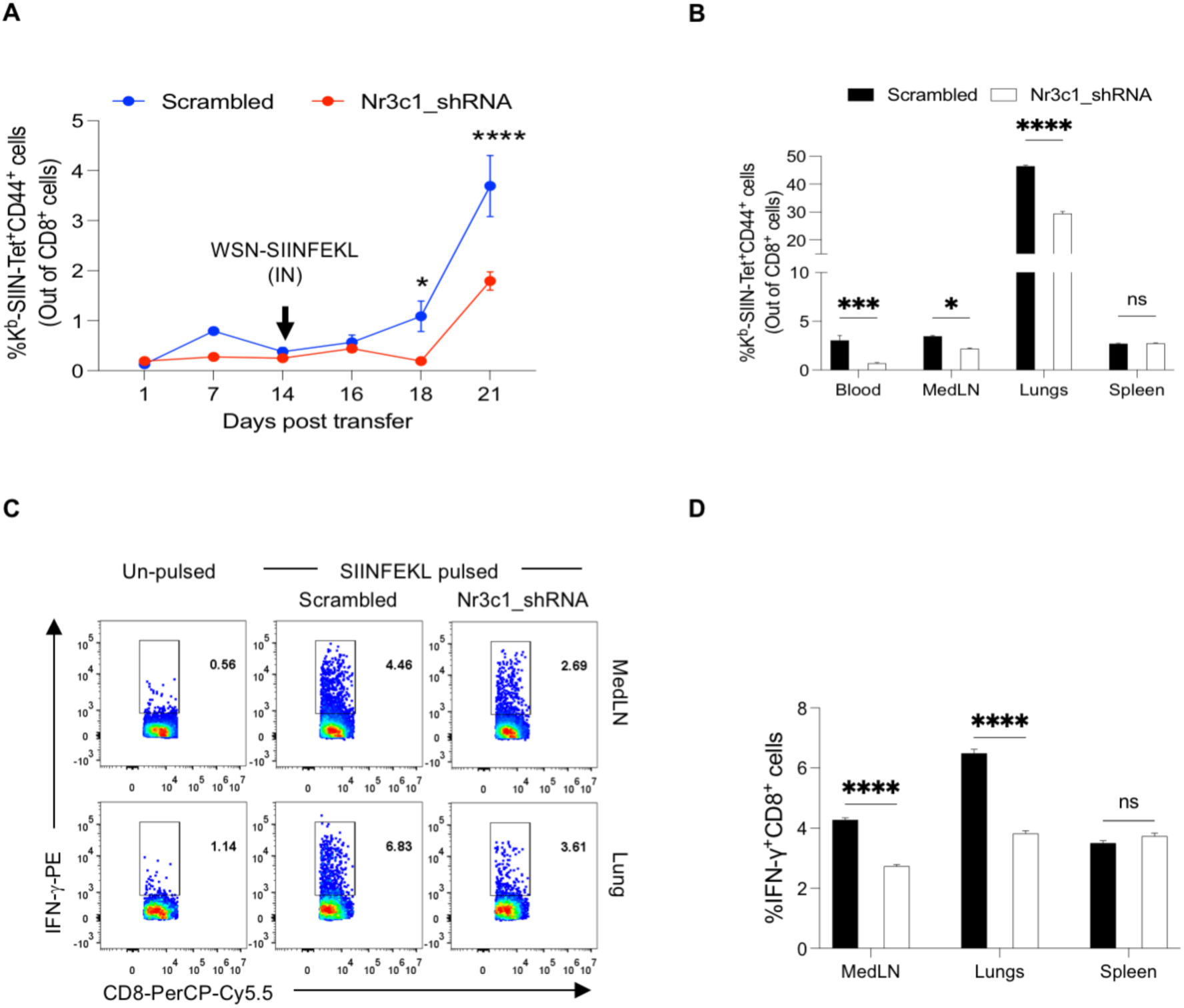
shRNA mediated knockdown of nr3c1 attenuates memory CD8^+^ T cell generation. OT1 cells were stimulated *in vitro* with anti-CD3 and anti-CD28 antibodies for 48 hours. These cells were then transduced with retroviruses, containing the shRNA for *nr3c1* or the scrambled sequence, by spinning at 800x*g* for 90 minutes at 37°C. 5 days after transduction, the cells were transferred into naïve sex matched B6 mice and were tracked for 2 weeks. 14 days after the transfer, the animals were infected intranasally with 200pfu of WSN-SIINFEKL. The recall kinetics in the two groups was measured. 7 days after the transfer, the animals were sacrificed and organs were collected for cellular analysis. **A.** Plot summarizes the kinetics of K^b^-SIINFEKL- Tet^+ve^ cells at different time points. Data represents Mean ± SEM; ***p<0.001; **p<0.005; *p<0.05 and ns (p>0.05)- not significant (Two-way ANOVA). **B.** Bar diagrams show the frequency of K^b^-SIINFEKL-Tet^+ve^ cells in the different compartments. Data represents Mean ± SEM; ***p<0.001; **p<0.005; *p<0.05 and ns (p>0.05)- not significant (Two-way ANOVA). **C, D.** Cytokine production by CD8^+^ T cells in lungs and medLN of each group was measured by ICCS assay. **C.** Representative FACS plots for the ICCS assay to measure IFN-γ production upon peptide stimulation are shown. **D.** Bar diagram summarizes the frequencies of CD8^+^ T cells producing IFN-γ upon peptide stimulation. Data represents Mean ± SEM; ***p<0.001; **p<0.005; *p<0.05 and ns (p>0.05)- not significant (Two-way ANOVA). The experiments were repeated two times and similar results were obtained with n=4 per group.

Next, we analyzed the transcriptome of polyclonal CD8^+^CD44^+^ cells sorted from sham and dexa treated MHV68 infected animals as this could offer analysis on T cell receptor (TCR) usage (Figure S6B). Genes with ≥ 1.4-fold change in FPKM values between Dexa T_x_ /Sham T_x_ in each of the gene sets represented pathway enrichment for oxidative phosphorylation (high expression), nucleocytoplasmic transport (intermediate expression), transport of small molecules, activation of matrix metalloproteinases, AMPK signaling, TCR signaling, JAK-STAT signaling, TGF-β signaling and cell cycle progression (low expression) (Figure 5A, Table 1). Since *Il6st* (∼2.7 fold), *Il6ra* (∼1.91 fold) and *Il7r* (∼2.4 fold) increased in the dexamethasone treated cells, we analyzed downstream signaling processes due to their role in driving the differentiation of memory CD8^+^ T cells (*2, 15*). The dexamethasone treated cells showed higher phosphorylation of STAT3 and STAT5β supporting for enhanced activities via these receptors (Figure 5D, E, and F). Interestingly, the dexamethasone treated cells preferentially expressed a shorter variant of STAT5β (∼70kDa) (Figure 5F, red arrowhead). Phosphorylation of the truncated but not full-length STAT5 was shown to be more stable and truncated STAT5 products have been shown to have enhanced nuclear localization (*16*). Such a switch might cause sustained signaling via IL6/IL7 receptors in the dexamethasone treated cells that led to their efficient memory differentiation.

**Figure 5.**
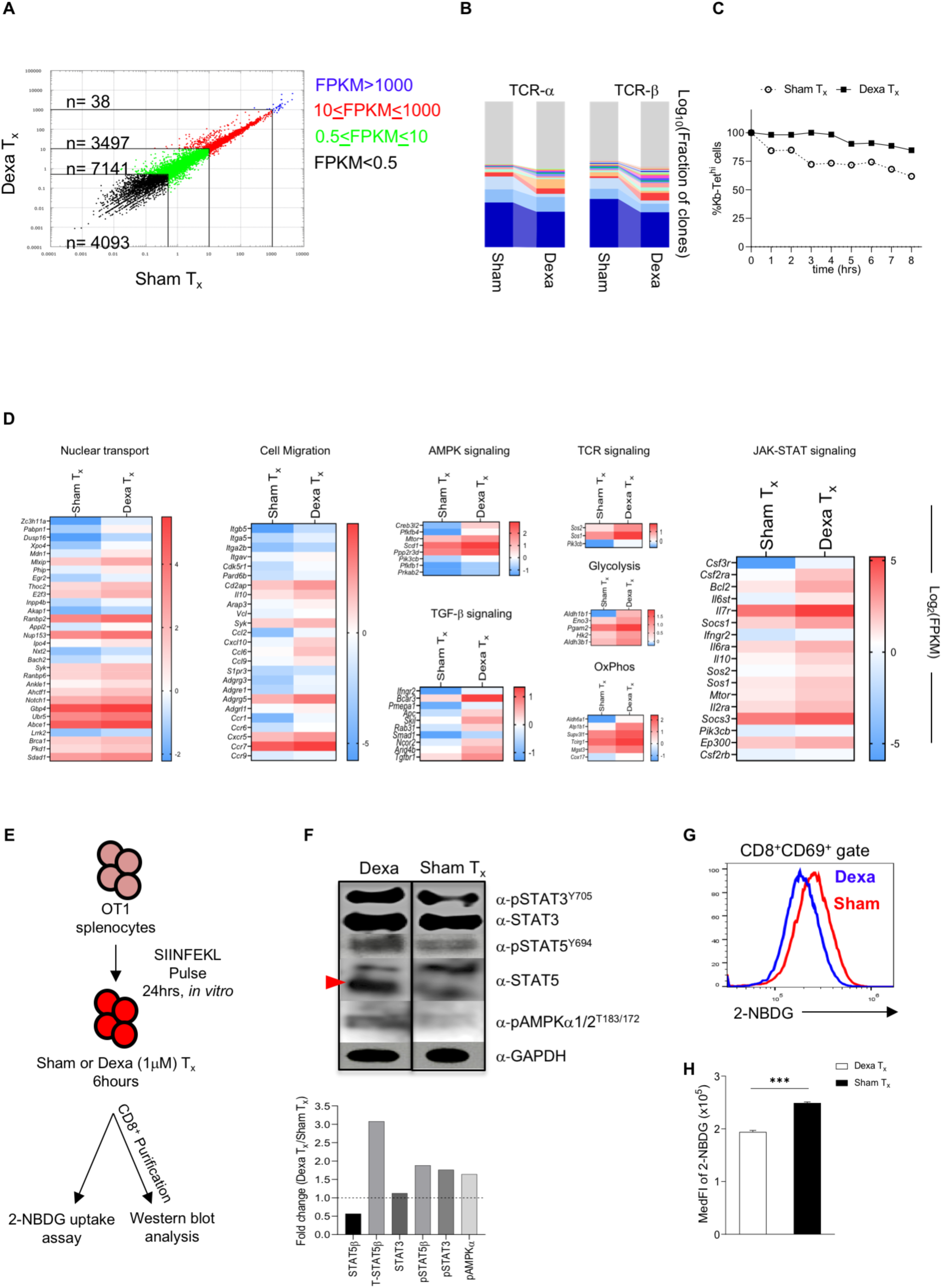
Mechanistic insights into glucocorticoid mediated memory CD8+ T cell differentiation. **A.** Scatter plot shows the differential expression of genes in Sham and Dexamethasone treated cells. **B.** TCR repertoire analysis was also performed. Stacked bar diagrams summarizing the relative abundance of mapped clones in the alpha and beta chain of Dexamethasone and Sham treated cells are shown. **C.** Plot summarizing the data obtained from tetramer dissociation assay is shown. **D.** Heatmaps for various genes involved in the mentioned pathways are shown. **E.** OT1 splenocytes were pulsed with SIINFEKL for 24 hrs, followed by dexamethasone or sham treatment for 6 hrs. Complete splenocytes were used for 2-NBDG uptake assay. In a separate experiment, the CD8^+^ T cells were also enriched by MACS and cell lysate was prepared for further analysis. **F.** Western blot analysis for the respective molecules is shown. Fold change calculate via densitometry is summarized in the bar diagram. **G.** Representative overlay histogram for the 2- NBDG uptake assay is shown. **H.** Bar diagram summarizing the analysis of 2-NBDG uptake assay is shown. Data represents Mean ± SEM; ***p<0.001; **p<0.005; *p<0.05 and ns (p>0.05)- not significant (Student’s T-test).

Genes involved in cell cycle progression such as *Cdk12* (∼1.9 fold)*, Ccnj* (∼2.3 fold)*, Ccnt2* (∼2.1 fold)*, Ccnl1* (∼1.4 fold) and *Ccnl2* (∼1.7 fold) were upregulated while the cyclin dependent kinase inhibitor 3, *Cdkn3* was downregulated by ∼1.4 fold. These results could explain elevated rates of cell division in the dexa treated cells. Moreover, the expression of the antiapoptotic gene, *Bcl2,* induced following JAK3/STAT5 axis activation (*12, 15*) was upregulated by ∼2-fold in the dexamethasone treated cells. Therefore, signaling via IL6 and IL7 receptors caused phosphorylation of STAT3 and STAT5 not only to promote cell division but also to induce a preferential survival of such cells.

AMPK signaling has been implicated in the initiation of fatty acid metabolism, a canonical switch from effector to memory CD8^+^ T cells (*17*). Moreover, upregulation of the genes encoding the mitochondrial enzymes such as *mt-Nd1* (∼1.6 fold)*, mt-Nd2* (∼1.71 fold)*, mt-Nd4* (∼1.5 fold)*, mt-Cytb* (∼1.6 fold), involved in oxidative phosphorylation in dexamethasone exposed cells further supported this shift (*18*). That the exposure to GCs induced low glucose metabolism in CD8^+^ T cells was shown previously (*19*). Therefore, an altered glucose metabolism could induce AMPK signaling (*20*). However, our transcriptome analysis revealed an upregulation of genes involved in glycolysis. Therefore, to reconcile such paradoxical observations, we explored the possibility of a compromised glucose uptake inducing AMPK signaling (Figure 5D, E and F). The dexamethasone treated cells showed a decreased uptake of glucose as compared to the sham treated cells when cultured in the presence of the fluorescent analog of glucose, 2-(N-(nitrobenz-2-oxa-1,3-diazol-4- yl) amino)-2 deoxy-D-glucose (2-NBDG) (Figure 5E and G). Therefore, an energy imbalance due to reduced glucose uptake could induce signaling via AMPK, a master regulator of ATP metabolism. Accordingly, the dexamethasone exposed cells as compared to the control cells expressed elevated level of phosphorylated AMPKα (Figure 5F). The induction of AMPK signaling could result in the inhibition of mTORC1 to promote memory CD8^+^ T cells generation (*21*). Therefore, GC signaling in the responding CD8^+^ T cells drove the differentiation of memory CD8^+^ T cells involving AMPK and JAK-STAT pathways. Several genes involved in TGF-β pathway were highly expressed in dexamethasone treated cells suggesting for the role of this pathway in imparting the differentiating cells a tissue residency phenotype (Figure 5D).

We also analyzed the TCR repertoire in each group and observed a preferential expansion of some clones in the dexamethasone treated cells as compared to the control cells (Figure 5B). The preferentially expanded and selected clones that formed lasting memory could have TCRs of higher affinity as T cells of low affinity have been shown to be susceptible to GC mediated apoptosis (*22*). Accordingly, at 7dpi, the virus-specific cells in the draining medLNs of dexamethasone treated animals had longer pMHC-TCR interactions as measured by tetramer dissociation assay (Figure 5C). Since our transcriptome revealed an unaltered expression of inhibitory molecules such as PD1 (*pdcd1*), TIM-3 (*havcr2*), LAG3 (*lag3*), GILZ (*Tsc22d3*), MKP1 (*Dusp1*), IκBα (*Nfkbia*) in the stimulated dexamethasone exposed cells, a regulated pMHC-TCR interaction in the dexamethasone treated cells with higher signal strength might have helped them retain functionality, particularly evident during the recall of the generated memory cells.

Next, we compared our transcriptome data with the gene expression profiles evident in naïve, effector and memory CD8^+^ T cells using the gene sets data available at (http://www.gsea-msigdb.org/gsea/msigdb/genesets.jsp?collection=IMMUNESIGDB). The dexamethasone treated cells exhibited a transcriptional profile more commonly observed in memory cells as well as those induced in response to IFN-γ, Type I IFN and TNF-α signaling (Figure S7A-D). Gene sets shown to be either upregulated or down-regulated in T_RM_ cells were largely represented in a similar pattern in the dexamethasone exposed cells (Figure S7E and F). This suggested for the functional superiority and enhanced T_RM_ transition of dexamethasone treated cells (*23*). To confirm the same, we administered the sham or dexamethasone treated mice in the late memory phase (>300dpi) of IAV infection with FTY720, an antagonist of sphingosine-1-phosphate receptor 1 (S1PR1) three days prior to re-infection. The treatment was continued until the termination of experiments (Figure S8A). In the draining medLNs, peripheral circulation, lungs and spleens of dexamethasone treated animals, frequencies of donor OT1 cells were upto two-fold higher but such cells were reduced in the distal inguinal LNs (Figure S8D). Despite the FTY720 induced sequestration of antigen reactive OT1 cells in LOs, more than two-fold higher frequencies of activated CD44^+^CD8^+^ T cells as well as donor cells were present in the infected lung tissues (Figure S8C). Interestingly, the mice injected previously with dexamethasone exhibited an increased proportion of virus-specific CD8^+^ T cells in the peripheral circulation, draining medLN and spleen indicating the elevated basal levels of memory pool in these organs. That a greater proportion of OT1 cells in medLN and spleen produced IFN-γ and TNF-α than those recovered from the lung tissues of the dexamethasone treated animals as compared to the control animals could suggest an efficient local control of the infection in the former group and consequently the cells were already in the contraction phase (Figure S8E-H).

Therefore, our transcriptional and biochemical analyses were further supported by our experimental data to show a better survival, migration and retention of antigen-specific CD8^+^ T cells at the site of infection. Enhanced levels of GC signaling in the responding cells induced AMPK signaling via a reduction in glucose uptake. Additionally, enhanced JAK-STAT3/5 signaling induced by a synergistic activity by of IL6 and IL7 supported the differentiation of memory CD8^+^ T cells with tissue residency.

Finally, we eliminated the possibility of dexamethasone influencing other cells to help fine tune the differentiation program of CD8^+^ T cells and stimulated OT1 cells *in vitro* with α-CD3 and α-CD28 antibodies. Thereafter, the cells were either exposed to dexamethasone (1μM) or the diluent for one hour. The adoptively transferred cells were analysed before or after infection with MHV68-SIINFEKL (Figure 6A). The cells were equally distributed in the peripheral circulation one day post transfer but their frequencies as well as numbers were more than two-fold higher in mice receiving dexamethasone treated 6 days later (Figure 6B & C). We observed significantly higher frequencies and numbers of K^b^-SIINFEKL-tetramer^+ve^ CD8^+^ T cells in bronchoalveolar lavage (BAL) and spleens of infected mice at 8dpi in the animals receiving Dexa T_x_ cells than those receiving the control cells (Figure 6D-G). Slight but significantly increased frequencies and numbers of K^b^-SIINFEKL-tetramer^+ve^ cells as well as those producing IFN-γ were also present in the medLNs of animals receiving dexamethasone treated cells (Figure 6F-H). The virus titers were ∼100-fold lower in the lung tissues of the animals receiving dexamethasone pulsed cells as compared to those injected with control cells (Figure 6J). A lesser degree of tissue disruption in lung parenchyma, lower cellular infiltrates and more alveolar spaces were clearly evident in animals receiving dexamethasone treated cells as compared to control cells suggesting for a better protection rendered by the dexamethasone cells during their recall (Figure 6K). The data showed that the dexamethasone pulsed antigen-specific CD8^+^ T cells showed better survival and provided protection against virus infection. Therefore, GC signaling in the responding CD8^+^ T cells induced a differentiation program to favor memory transition of effector cells and such cells had higher propensity to populate peripheral sites of infection. Such an approach could therefore be useful in generating a better memory T cell repertoire.

**Figure 6.**
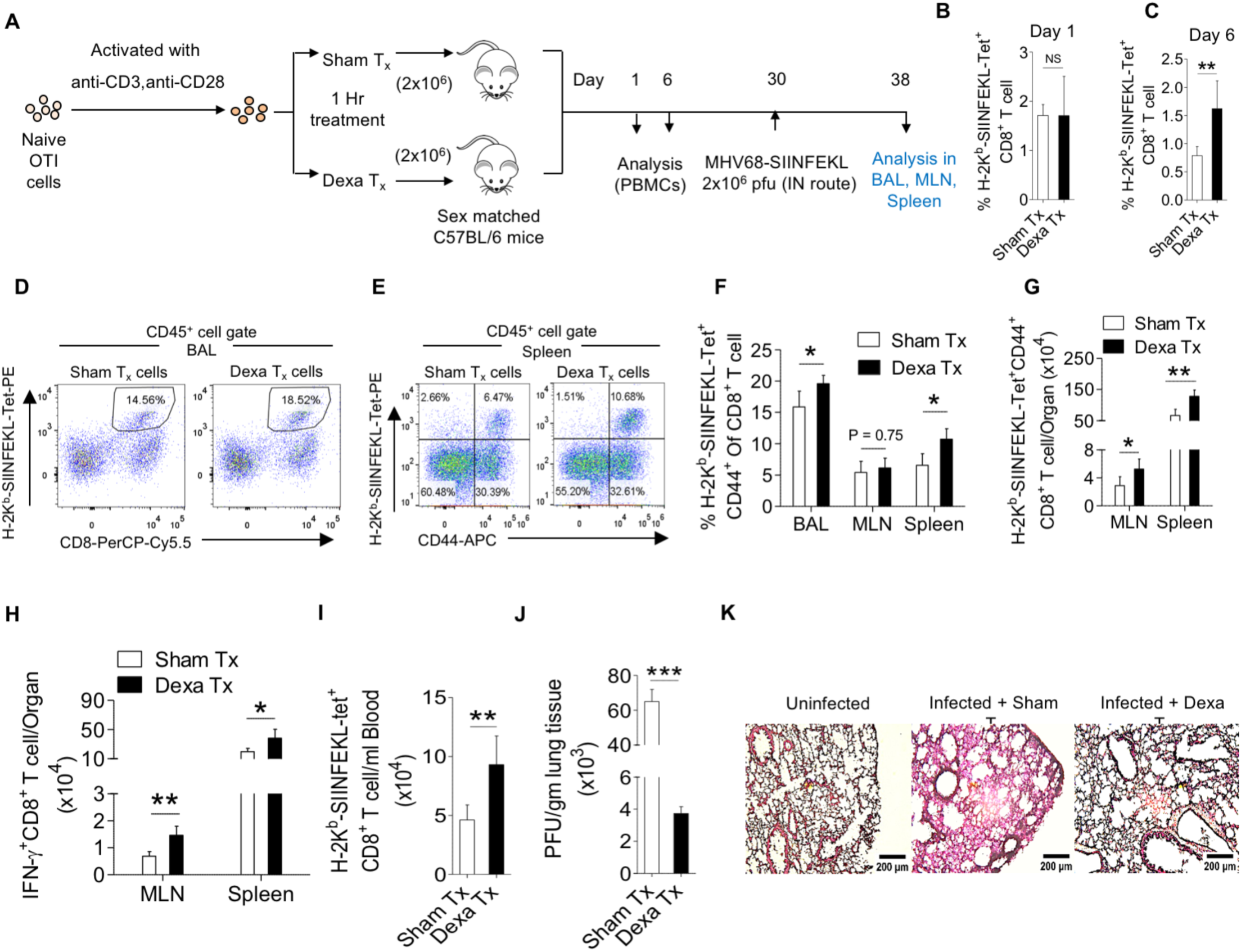
Improved memory transition and recall of GC exposed activated CD8^+^ T cells. **A.** OT1 cells were *in vitro* stimulated with anti-CD3 and CD28 antibodies for 16 hours. These cells were then exposed for one hour either to dexamethasone (1μM) or to the diluent only. 2×10^6^ cells per mouse were adoptively transferred separately in sex-matched mice and their fates were tracked in the recipient animals subsequently. **B, C**. Bar diagrams summarize the frequency of tetramer positive cells in the circulation of recipients 1 day (**B**) and 6 days (**C**) after adoptive transfer. Data represents Mean ± SD; ***p<0.001; **p<0.005; *p<0.05 and ns (p>0.05)- not significant (Unpaired t test). **D, E**. Representative FACS plots show K^b^-SIINFEKL-tetramer^+ve^ cells in the bronchioalveolar lavage (**D**) and spleen (**E**) of each group. **F, G**. Bar diagrams summarize the frequency (**F**) and count (**G**) of K^b^-SIINFEKL-tetramer^+ve^ cells in the various organs analyzed. Data represents Mean ± SD; ***p<0.001; **p<0.005; *p<0.05 and ns (p>0.05)- not significant (Unpaired t test). **H**. Bar diagram summarizing the count of IFN-γ producers in the organs analyzed is shown. **I**. Bar diagram summarizing the frequency of K^b^-SIINFEKL-tetramer^+ve^ cells in circulation of each group is shown. **J**. Bar diagram summarizing the results of plaque assay to measure viral titers in lungs of each group is shown. Data represents Mean ± SD; ***p<0.001; **p<0.005; *p<0.05 and ns (p>0.05)- not significant (Unpaired t test). **K**. Representative sections of lung tissues stained by H&E are shown as histological micrographs from different groups of animals. Scale bar=200μm. The experiments were repeated two times with similar results.

This study demonstrated the role of glucocorticoid signaling in the differentiation of functional, tissue-homing CD8^+^ T cell memory. That a transient dexamethasone therapy in the expansion phase of T cells during a primary infection augments long-term memory with tissue homing potential was shown. Blocking endogenous GC responses with a competitive inhibitor of the GR generated SLECs while a transient therapy with a long-acting synthetic analog of GCs, dexamethasone, preferentially generated MPECs that further differentiated into long-term tissue residing functional memory cells. By expanding locally in lung tissues following a secondary infection, such cells provided efficient viral control. The cell intrinsic role of GC signaling on the memory transition of effector cells was shown by genetic depletion of the GR in the antigen- specific cells. Transcriptome and biochemical analysis revealed a GC mediated enhancement of T_RM_ response by an engagement of AMPK, JAK/STAT and TGF-β signaling pathways. We also described an *ex vivo* strategy to efficiently drive the differentiation of stimulated CD8^+^ T cells into memory cells using dexamethasone. Such a therapy could add value to cell-based immunotherapies aimed at managing chronic infection and cancers. However, a sustained high level of GC signaling in the tumor microenvironment can result in the exhaustion of effector CD8^+^ T cells (*24*). Therefore, a transient exposure is critical for the enhancement of memory response. Whether or not GCs should be used to suppress immunopathological reactions caused by viruses is debated as the compromised immunity could increase virus loads, thereby inducing dysregulated responses. Therefore, a transient GCs therapy could be considered when the infectious agent is adequately controlled. Our observations could be directly relevant and tested in COVID-19 patients who recovered following a dexamethasone therapy by analyzing the functionality of SARS-CoV2 specific CD8^+^ T cells in the PBMCs. Furthermore, the reinfection rates and the resolution of infection in such individuals could provide a valuable clinical correlate of dexamethasone promoted local protective response. In addition to adding another layer to the complicated process of T cell differentiation, we also provide a primer for future investigations aimed at elucidating the long-term implications of corticosteroid therapy in fine tuning immunological memory during viral infections.

## Acknowledgments

We thank Dr. Rajesh Ramachandran’s group at IISER, Mohali for the help provided in confocal microscopy. The help provided by the flow cytometry facility at IISER, Mohali is appreciated.

## Author contributions

Conceptualization: S. Sehrawat, A.T. & D.K.; Methodology: S. Sehrawat, A.T., D.K., R.S., S.S., A.D.; Writing- Original draft: S. Sehrawat, A.T.; Writing- Review and Editing: S. Sehrawat, A.T.; Visualization & Data Presentation: S. Sehrawat, A.T.; Data curation & formal analysis: A.T., D.K., R.S., S.S., A.D.; Supervision: S. Sehrawat; Funding acquisition: S. Sehrawat.

## Supplementary Materials

### Materials and Methods

#### Mice and viruses

C57BL/6 (Stock No. 000664), B6 OT1 (C57BL/6-Tg(TcraTcrb)1100Mjb/J; Stock No. 003831), and B6 CD45.1 (B6.SJL-Ptprc^a^ Pepc^b^/BoyJ; Stock No. 002014) were procured from Jackson Laboratory, USA. All the animals were housed and bred in the individual ventilated cages in the small animal facility for experimentation (SAFE) of the Indian Institute of Science Education and Research (IISER), Mohali. The animal experiments were performed strictly in accordance with the protocol approved by the Institutional Animal Ethics Committee (IAEC), IISER Mohali, constituted under the aegis of Committee for the Purpose of Control and Supervision of Experiments on Animals (CPCSEA). MHV68-SIINFEKL and IAV (WSN-SIINFEKL) were used for *in vivo* experiments. HSV1 MHV68-SIINFEKL was propagated, harvested and titrated using Vero cells and stored at -80°C until further use as described earlier. WSN-SIINFEKL was propagated and titrated using MDCK cells. WSN-SIINFEKL was used for intranasal infections at the indicated doses. MHV68-SIINFEKL was used for infecting intraperitoneally or inducing heterologous challenge intranasally in mice previously infected with WSN-SIINFEKL at the indicated doses.

#### Viral infection and drugs administration in mice

For the adoptive transfer experiments, OT1xRAG1^-/-^ mice were sacrificed and CD8^+^ T cells were MACS purified from single cell suspensions of the pooled lymph nodes and splenocytes. Specific number of cells were then transferred into CD45.1^+^ congenic mice. For all the adoptive transfer experiments, donor and the recipient animals were sex matched and of the same age group. The frequency of the donor cells in circulation was checked just before infection with WSN-SIINFEKL via the intranasal (i.n) route. The animals were then given recurrent doses of dexamethasone (10 mg/kg body weight) from 4 to 6 dpi or mifepristone (40 mg/kg body weight) from 1 to 5 dpi or the diluent. The donor OT1 cells were tracked in circulation. A heterologous challenge was given i.n with MHV68-SIINFEKL after 3, 4 or 12 months post primary infection and the recall response was monitored by analyzing the frequency of OT1 cells in circulation at different days post re- infection (dpr). The animals were sacrificed at indicated time following re-infection from each of the groups and the collected organs were subjected to cellular analysis. Similar experiments were performed for measuring the optimal dose response of dexamethasone and mifepristone administration.

*In vitro* stimulation of OT1 cells with α-CD3 and α-CD28 was performed with plate-coated antibodies for 16 hours followed by their exposure to 1μM dexamethasone or the diluent for 1 hour. After extensive washings, the cells were transferred in separate groups of animals and their survival and retention was measured at one and six days after transfer. The transferred cells were recalled by infecting recipients intranasally with MHV68-SIINFEKL (2×10^6^ pfu) after 30 days of transfer. The cells were analyzed at indicated time points by flow cytometry.

#### Antibodies, and other biological reagents

Antibodies used for measuring the expression of different molecules were procured from BD Biosciences, Tonbo biosciences, eBiosciences and BioLegend. The antibodies used were against CD8-PerCP Cy5.5 (530-6.7), CD45.2-APC, -FITC & -PE (1O4), CXCR3-FITC (173), CD44-APC (IM7), CD62L-APC (MEL 14), CD69-APC (H1.2F3), CD127-APC (A7R34), KLRG1-AF488 (2F1), CD103-FITC (2E7), IFN-γ-PE (XMG1.2), TNF-α-APC (TN3-19), Granzyme B-PE (NGZB). The MHC class I tetramers used in this study were synthesized in house and the monomers were refolded with SIINFEKL or SSIEFARL. All the antibodies were diluted in FACS buffer (Phosphate Buffered Saline (PBS) with 2% FBS) in 1:200 ratios. Dexamethasone, mifepristone, haematoxylin, eosin Y and 2,2,2- tri-bromoethanol were purchased from Sigma Aldrich. MojoSort CD8^+^ T cell purification kit was purchased from BioLegend. CFSE, intracellular fixation-permeabilization buffer, and Streptavidin-PE were purchased from Thermofisher. Monensin, Brefeldin A, purified anti-CD3ε (17A2) and anti-CD28 (37.51) were purchased from eBioscience. The primary antibodies for western blot analysis were against STAT3, phosphoSTAT3(Y705), STAT5β, phosphoSTAT5β(Y694) and phospoAMPKα1/2(T183/172). The PhosphoPair STAT3 (Tyr705) Antibody Set was purchased from Biolegend (Cat. 699952) and the primary antibodies against STAT5β (Cat. 71-2500), phosphoSTAT5β(Y694) (ST5P-4A9, Cat. 33-6000) and phospoAMPKα1/2(T183/172) (Cat. PA5-17831) were purchased from Invitrogen. GAPDH loading control antibody was purchased from Invitrogen (GA1R, Cat. MA5-15738). 2-NBDG was purchased from Invitrogen.

#### Flow cytometry for cellular analysis

For blood staining, blood samples were collected in EDTA tubes. 25μl of blood was aliquoted directly in a centrifuge tube and 5μl of antibody mix was added. The cells were then incubated at 4°C for 45 minutes in dark. After incubation, 300μl of 1x ACK solution (155mM NH_4_Cl, 12mM NaHCO_3_, 0.1mM EDTA, pH 7.3) was added to lyse RBCs. The cocktail was kept at room temperature for 10 minutes followed by addition of 300 μl of PBS. The cells were then acquired and analyzed by flow cytometry using BD C6 flow cytometer or BD FACSAria Fusion.

The mice were sacrificed and perfused with 25ml of PBS via the left, then right ventricle to remove residual circulating cells from organs by incising inferior vena cava traversing through diaphragm in the abdominal cavity. Different lymphoid and non-lymphoid organs were collected from the mice. Single cell suspensions were then prepared. In brief, the organs were placed in a 70μm cell strainer with 2ml of cold complete RPMI and gently crushed using the soft end of a 2.5ml syringe plunger. All the cells were passed through the strainer and collected in a 15ml centrifuge tube. The cells were washed twice with cold PBS by centrifugation at 1200rpm for 5 minutes at 4°C. The cells were finally resuspended in complete RPMI for further cellular analysis. The non-lymphoid organs were prepared by first digesting with type IV collagenase for 1 hour at 37°C. Single cell suspensions were then made as described above. The cells were stained using the indicated fluorescent antibodies at 4°C for 30 minutes. For intracellular staining, cells were first surface stained and then treated with fixation buffer (eBiosciences) for 20 minutes. This was followed by permeabilization using intracellular permeabilization buffer from eBiosciences. The stained cells were washed three times with cold PBS and acquired using BD Accuri C6 or BD FACS Aria fusion. FlowJo v10 software was used for analysis of the acquired data.

For the 2-NBDG uptake assay, dexa or sham treated activated cells were incubated with 10μM of 2-NBDG for 30 minutes at 37°. The cells were immediately placed on ice and washed five times with cold PBS. The washed cells were then stained with α-CD8 and α-CD69 antibodies and incubated on ice for 1 hour. The cells were washed twice with cold PBS and acquired using BD Acuri C6 flow cytometer. FlowJo v10 software was used for analysis of the acquired data.

#### Tetramer Dissociation Assay

Cells isolated from the draining mediastinal lymph node of WSN-SIINFEKL infected mice that were sham or dexamethasone treated. These cells were stained with an excess of respective tetramers and α-CD8 antibody for 30 min at 4°C and washed three times with cold PBS. The stained cells were then incubated at 16°C for varying periods of time. Aliquots were removed at indicated time points from the suspended cells and fixed immediately with 1% PFA in FACS buffer. At the end of the experiment samples were analysed flow cytometrically.

#### Histopathology and Confocal microscopy

The caudal right lobe of lungs from each group of mice were collected and fixed overnight at 4°C in 4% PFA prepared in 10 mM PBS. Tissues were then dehydrated at room temperature in a 5%- 20% gradient of sucrose prepared in 10mM PBS at 4°C . The tissues were embedded in OCT compound and the tissue blocks were used for cutting tissue sections of 5µM on a Leica Cryotome. The sections were stained by H & E. The dried sections were imaged using a microscope from Leica (DMi8). The images were analysed using ImageJ software.

For confocal imaging, the sections were prepared as mentioned above. They were then washed 3 times with 1x 0.1% PBST (0.1% Triton-x100 in 1x PBS). After blocking with 1% BSA prepared in 1x PBST for 1 hour, the slides were washed twice with 1x PBST and left overnight for staining at 4°C. After 2 washes with 1x PBST, they were stained with Hoechst for 1 minute, followed by three washes with 1x PBST. They were then given a final wash with autoclaved MilliQ water. The stained sections were visualised using Olympus FV10 confocal laser scanning microscope at the indicated magnification.

#### Quantification of MHV-68-SIINFEKL in lung tissues

The mice were sacrificed at the indicated time post infection and the lung tissues were collected from PBS perfused mice infected with MHV68-SIINFEKL. The organs were weighed and homogenates of lung tissues were prepared to determine viral burden on Vero cells.

#### Lentivirus based RNAi

Double stranded oligonucleotides for short hairpin RNA (shRNA) against *nr3c1* were cloned into pLKO.1-GFP between AgeI and EcoRI restriction enzyme sites. The sequences for the shRNA are: 5’-CCGGACCGGTCAGGCTGGCTTTATTAAATTCAAGAGATTTAATAAAGCCAGCCTGGAATTCC-3’ (top strand) and 5’- GGAATTCCAGGCTGGCTTTATTAAATCTCTTGAATTTAATAAAGCCAGCCTGACCGGTCCGG-3’ (bottom strand). Vector specific primers were used to screen positive colonies by PCR. The recombinant lentivirus was made by co-transfection with pLKO.1GFP-nr3c1shRNA, pCMVR8.74 (Packaging vector), pMD2.G (Envelope vector), Tat and Rev Plasmids in HEK293T cells using polyethyleneimine (PEI, stock conc: 1mg/ml). For transfection of 100mm dish with 70- 80% confluency, the concentrations of the aforementioned were as follows: pLKO.1GFP- nr3c1shRNA (10μg), pCMVR8.74 (9μg), pMD2.G (6μg), Tat (6μg) and Rev plasmid (6μg). These were mixed in 1ml of serum free DMEM along with 1/3^rd^ the volume of PEI (in μl). Mixing of all the components was done by vortexing the mixture for 30 seconds followed by incubation at room temperature for 15 minutes. This mixture was used for transfection of HEK293T. The medium was replaced with complete DMEM after 6 hours to remove extra PEI. 72 hours after transfection, the culture supernatants were collected and concentrated using 20% PEG-NaCl. The concentrated virus was then used for transduction of OT1 cells. A similar process was used for generating retroviruses containing the scrambled shRNA sequences. The sequences for the scrambled shRNA were as follows: 5’- CCGGACCGGTAGTATGGTTCAGATCGAGTTCAAGAGACTCGATCTGAACCATACTGAATTCC-3’ (top strand) and 5’- GGAATTCAGTATGGTTCAGATCGAGTCTCTTGAACTCGATCTGAACCATACTACCGGTCCGG-3’ (bottom strand). The knockdown was confirmed at a protein level by western blotting using anti-Nr3c1 Ab (Thermo Fisher Scientific, catalog no. PA1-512) (Figure S5A)

#### Lentivirus transduction for knockdown of *nr3c1* and RT-qPCR and *in vivo* analysis of GR lacking CD8^+^ T cells

To transduce OT1 cells, MACS purified OT1 cells were activated *in vitro* with α-CD3 and α- CD28 antibodies for 48 hours. These cells were then spin-transduced at 800 x*g* for 90 minutes at 37°C with freshly collected retrovirus. The cells were then kept in humidified CO_2_ incubator for 4 days. Total RNA was isolated from the nr3c1-shRNA and scrambled-shRNA transduced cells respectively by trizol method. The isolated RNA was then converted to cDNA using first strand cDNA synthesis kit (Verso cDNA synthesis kit, Thermo Fischer Scientific) according to the manufacturer’s protocol. Qualitative real time PCR (qPCR) was then performed using 2x-DyNamo ColorFlash SYBR Green qPCR kit from Thermofisher (F416L). The reaction was carried out using QuantStudio Real Time PCR system (ThermoFisher). The expression of HPRT gene served as the endogenous control and the relative expression of the various genes was calculated by 2^-ΔΔCT^ values. The reaction conditions used were: Initial denaturation (95°C for 7 minutes), denaturation (95°C for 10 seconds), annealing and extension (60°C for 30 seconds) for 40 cycles. Subsequently, melt curve analysis was performed. The primers used and the product size for the different genes were: nr3c1 (FP: 5′-AGTGATTGCCGCAGTGAAAT-3′ & RP: 5′- GCCATGAGAAACATCCATGA-3′) = 105 bp; Tbet (FP: 5′ CAATGTGACCCAGATGATCG 3′ & RP: 5′ GCGTTCTGGTAGGCAGTCAC 3′) = 168 bp; IFN-γ (FP: 5’- TGAATGTCCAACGCAAAGCA-3’ & RP: 5’-CTGGGATGCTCTTCGACCTC -3’) = 122; CD127 (FP: 5’-CAGCAAGGGGTGAAAGCAAC-3’ & RP: 5’- CTCGCTCCAGAAGCCTTTGA-3’) = 149 ; CD103 (FP 5’-CCACAGGACGAAGATCACTGT- 3’ & RP: 5’-CCCTCCTTGTGCTCTCCAAG-3’) = 169.

#### RNA sequencing and bioinformatic analysis

CD8^+^CD44^+^ cells were sorted at 8dpi from the spleens of MHV68-SIINFEKL infected mice that had received either three doses of dexamethasone from 4-6 dpi or the diluent. Samples from 12 mice were pooled for the analysis. RNA was isolated from the sorted cells by Trizol method. The qualities and quantities of the RNA samples were checked on NanoDrop followed by Agilent TapeStation using High Sensitivity RNA ScreenTape. The RNA-Seq paired end sequencing libraries were prepared from the QC passed RNA samples using illumina TruSeq Stranded mRNA sample Prep kit as per the manufacturer’s protocol. Briefly, mRNA was enriched from the total RNA using poly-T attached magnetic beads. Thereafter, enzymatic fragmentation was performed. First strand cDNA conversion was achieved using SuperScript II and Act-D followed by second strand synthesis. The double stranded cDNA was then purified using AMPure XP beads which was then followed by poly A-tailing and adapter ligation. A total of 15 PCR cycles were performed for enrichment. The PCR enriched libraries were purified using AMPure XP beads and analyzed on 4200 TapeStation system (Agilent Technologies) using high sensitivity D1000 Screen tape as per manufacturer instructions. After obtaining the Qubit concentration for the libraries and the mean peak sizes from Agilent TapeStation profile, the PE illumina libraries were loaded onto NextSeq500 for cluster generation and sequencing. The kit reagents were used in binding of samples to complementary adapter oligos on paired-end flow cell. The adapters were designed to allow selective cleavage of the forward strands after re-synthesis of the reverse strand during sequencing. The copied reverse strand was used to sequence from the opposite end of the fragment. Raw fastq files were assessed for quality using FASTQC and the 5’ end reads were trimmed accordingly using an in-house python script. The trimmed fastq files were subsequently mapped to the reference genome assembly mm10 of *Mus musculus* using Hi-SAT2. The output Bam files were processed using sam-tools and the processed bam files were used to count number of reads that map to each gene in the mm10 assembly using HT-Seq count. The raw read counts were converted to FPKM values using in-house python script. An initial cutoff was set on the number of raw reads obtained for each gene. Only the genes with reads ≥ 10 were kept for calculation of FPKM values. Genes having an FPKM ≥ 0.5 were then used for further analysis. The genes were classified into three groups based upon their FPKM values, *viz*, 0.5≤FPKM≤10 (Low expression); 10≤FPKM≤1000 (Intermediate expression); and FPKM>1000 (Highly expressed). The TCR repertoire analysis was performed using MiXCR.

#### Western blotting

OT1 splenocytes were pulsed with SIINFEKL for 24 hours. The cells were divided into two groups. One group was treated with 1μM dexamethasone while the other group was sham treated. 6 hours post treatment, the cells were washed thrice with cold PBS and lysate was prepared by using a Hypotonic Lysis Buffer, (HEPES 20 mM, EDTA 0.2 mM, MgCl2 1.5 mM,, KCl 100mM, 20%(V/V) Glycerol, 0.02%(V/V) NP-40, pH 7.5). The lysate was size fractionated in 12% denaturing acrylamide gel before transferring onto immune-blot PVDF membrane (Biorad, 162- 0177), followed by probing with the specific primary antibodies and HRP-Conjugated secondary antibody using Clarity Western ECL (Biorad, 170-5061). When necessary, the blots were stripped using an acidic stripping buffer ( For 1L buffer, 2.8g Glycine, 10g SDS, pH 2.2).

#### Statistical analysis

GraphPad Prism 9 was used for statistical analysis. Data with similar variances and having Gaussian distribution were analyzed with two tailed unpaired Student’s t-tests. Data not following Gaussian distribution were analyzed with two tailed Mann-Whitney U tests. For multiple comparisons, two-way ANOVA test with Bonferroni post hoc analysis were used. The p value below 0.05 were considered significant and were indicated as * <0.05, ** <0.01, *** <0.001 and ****<0.0001.

### Data and Code availability

The raw read counts and FPKM values for the transcriptome analysis are included in a file uploaded with the supplementary materials. Any additional information required to reanalyze the data reported in this paper is available from the corresponding author upon request. The python scripts used for analysis of transcriptome data are available at the following link- https://gist.github.com/dub2s/a8da867516d81dd1ed0b4c425c002722

## Supplementary Figures

**Figure S1.**
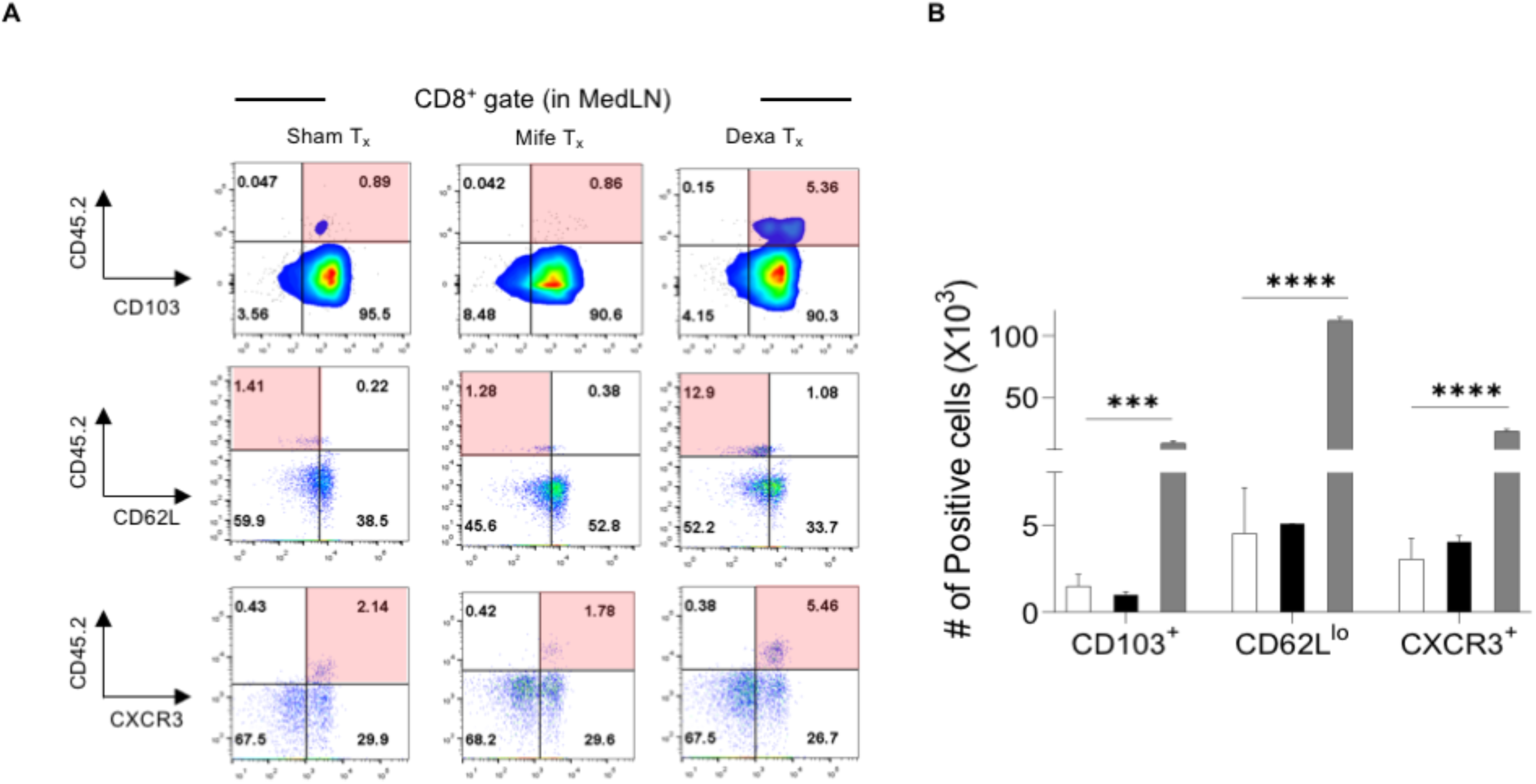
Phenotype of GC enhanced memory CD8^+^ T cells. A. Representative FACS plots show the frequency of cells expressing the indicated surface markers in the donor cells in the draining medLN. B. Bar diagram summarizing the total count of cells staining for the respective markers is shown. Data represents Mean ± SEM; ***p<0.001; **p<0.005; *p<0.05 and ns (p>0.05)- not significant (Two-way ANOVA).

**Figure S2.**
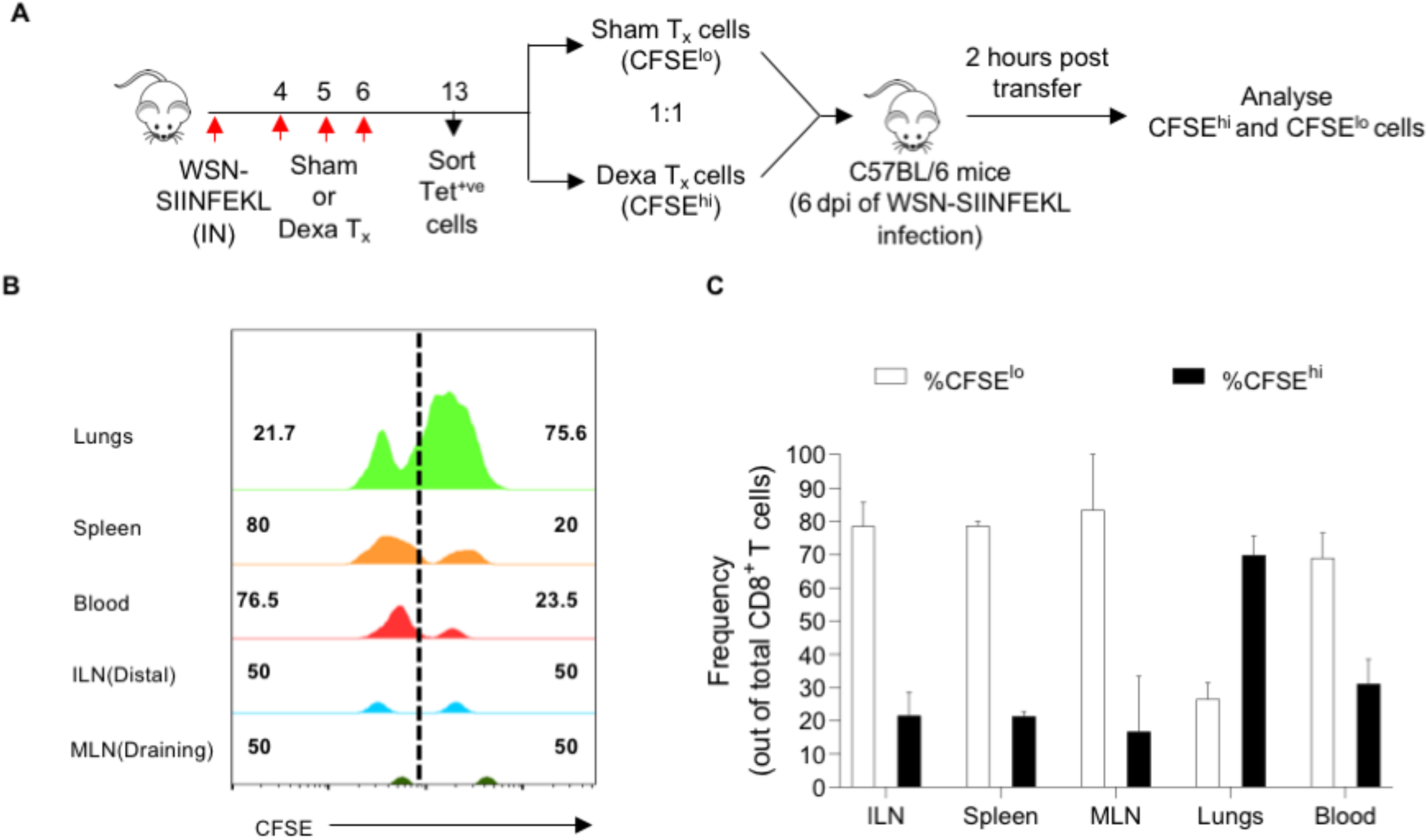
Migratory potential of GC exposed cytotoxic CD8^+^ T cells. A. A schematic of the experiment is shown. Animals infected with 500 pfu WSN-SIINFEKL via intranasal route were either given the diluent (Sham) or 10 mg/kg BWt. Dexamethasone from 4-6 dpi. K^b^-SIINFEKL-tetramer^+ve^ cells from each group were sorted at 13dpi from the draining medLNs. K^b^-SIINFEKL-tetramer^+ve^ cells obtained from the dexamethasone treated animals were labelled with a high concentration of CFSE (2.5mM; Dexa cells, CFSE^hi^) while cells from the sham treated animals were labelled with a ten-fold lower concentration of CFSE (0.25mM; Sham cells, CFSE^lo^). The labelled cells were then mixed in a 1:1 ratio and adoptively transferred into sex-matched mice that were in the acute phase of WSN-SIINFEKL infection. 2 hours after the transfer, the animals were sacrificed and organs were collected for analysis of the migration pattern of the transferred cells. B. Histograms show the relative distribution of CFSE^hi^ and CFSE^lo^ cells in the respective organs. C. Bar diagram summarizing the analysis done in various organs is shown.

**Figure S3.**
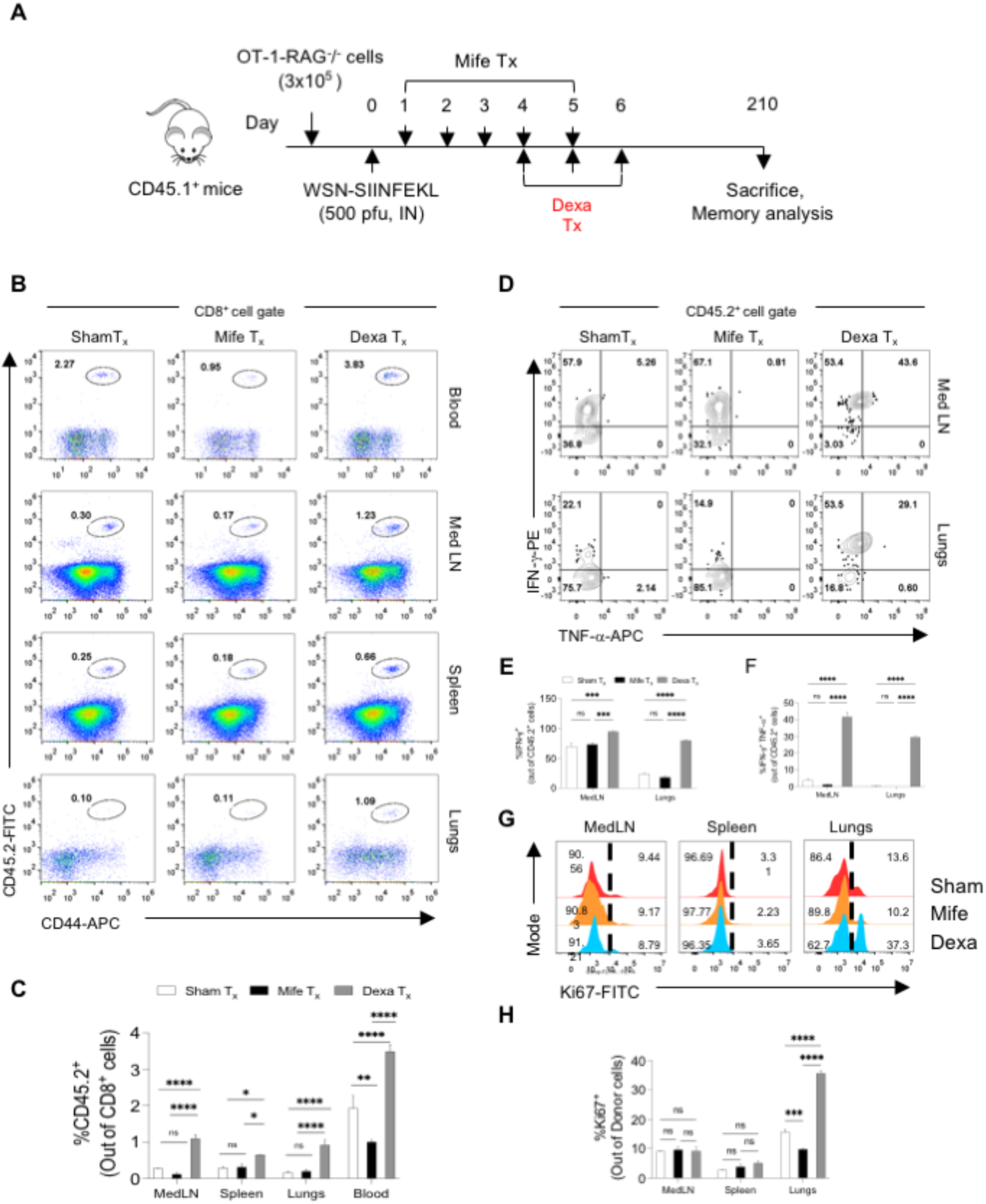
A transient dexamethasone therapy induces sustained virus-specific CD8^+^ T cell memory responses. A. A schematic of the experiments is shown. 3×10^5^ enriched OT-1 cells were adoptively transferred into sex-matched naïve congenic CD45.1^+^ B6 mice. The recipients were then intranasally infected with 500 pfu of WSN-SIINFEKL and injected with mifepristone, dexamethasone or diluent at the indicated timepoints. The lymphoid and non-lymphoid organs were then analyzed at 210 dpi for measuring long-term memory response of CD45.2^+^ donor cells. B. Representative FACS plots show the frequencies of donor cells in blood, medLN, spleen and lungs. C. Bar diagrams summarize the frequencies of donor cells in medLN, spleen, lungs and blood of each group. D. Representative FACS plots show the frequency of SIINFEKL stimulated IFN-γ and TNF-α producing donor cells in the medLN and lungs. E-F. Bar diagrams show the frequency of donor cells producing IFN-γ (E) and IFN-γ and TNF-α (F). G. Representative half-offset histograms for the analysis of Ki-67 positive donor cells in the medLN, spleen and lungs are shown. H. Bar diagrams summarizing the frequency of donor cells stained for Ki-67 in the indicated organs are shown. Data represents Mean ± SEM; ***p<0.001; **p<0.005; *p<0.05 and ns (p>0.05)- not significant (Two-way ANOVA).

**Figure S4.**
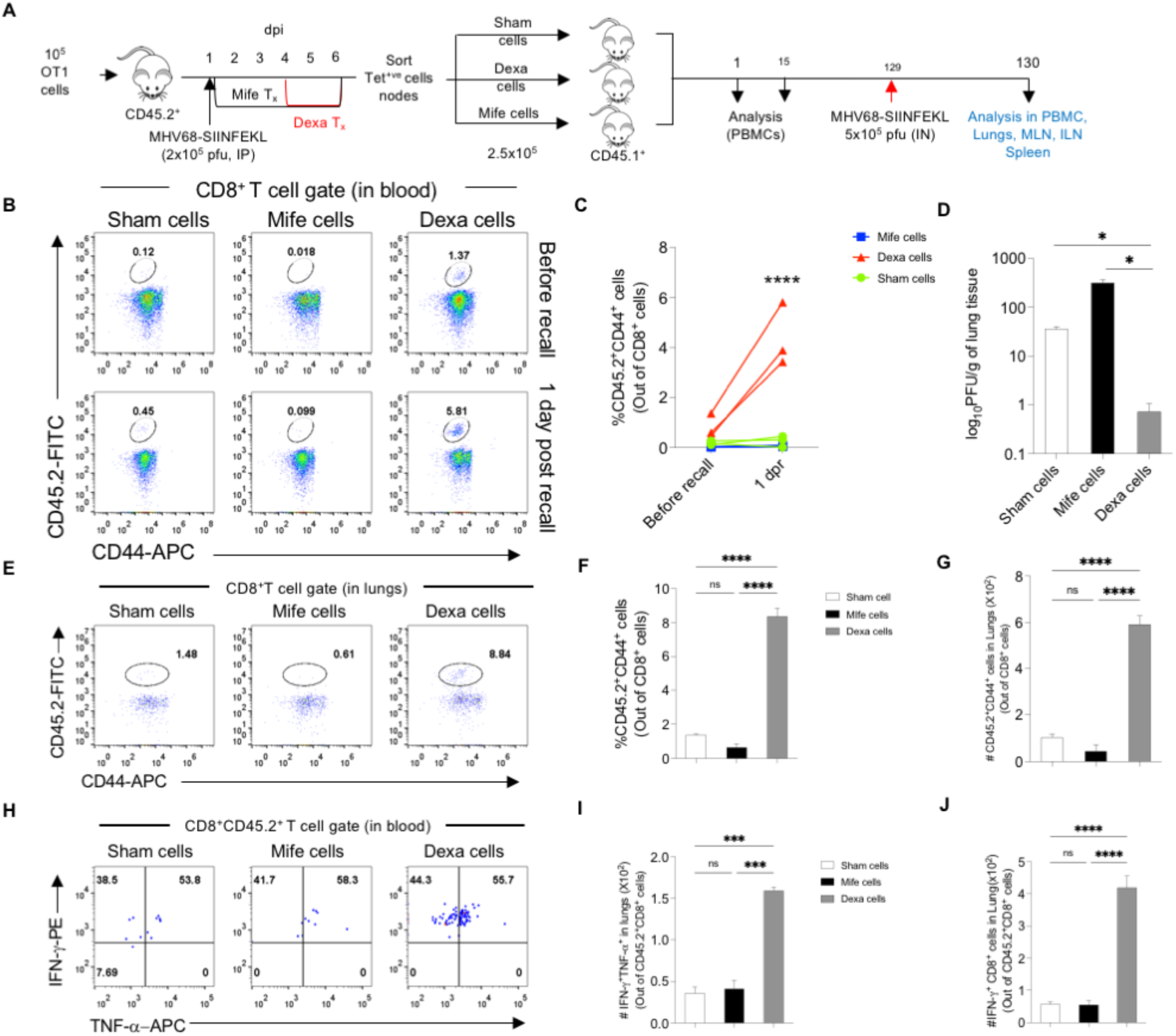
Enhanced survivability, recallability and functionality of GC exposed cytotoxic T cells. A. A schematic representation of the experiment is shown. 10^5^ naïve OT1 cells were transferred into B6 mice which were subsequently infected with 2×10^5^ pfu of MHV68-SIINFEKL (IP). This was followed by mifepristone, dexamethasone and sham treatment at the indicated timepoints. The mice were sacrificed at 7dpi and the tetramer positive cells were sorted from pooled lymph nodes of each group. The sham, mife and dexa exposed cells were then transferred into sex matched congenic CD45.1^+^ mice. The mice were then infected with 5×10^5^ pfu of MHV68-SIINFEKL at 129 days post transfer and the recall of the donor cells was tracked in circulation. The mice were sacrificed 1 day after infection and subjected to cellular analysis in various compartments. B. Representative FACS plots show the frequencies of CD45.2^+^ cells in circulation of each group. C. Plot summarizing the frequency of CD45.2^+^ cells in the circulation of the recipients before and 1 day after recall is shown. Data represents Mean ± SEM; ***p<0.001; **p<0.005; *p<0.05 and ns (p>0.05)- not significant (Student’s T-test). C. Bar diagrams summarize the viral titers obtained from the homogenised lung tissues of each group. Data represents Mean ± SEM; ***p<0.001; **p<0.005; *p<0.05 and ns (p>0.05)- not significant (One-way ANOVA). E. Representative FACS plots show the frequencies of CD45.2^+^ cells in the lungs of the recipients. F-G. Bar diagrams summarize the frequency (F) and absolute counts (G) of the CD45.2^+^ cells in the lungs of each group. H. Representative FACS plots show the proportion of CD45.2^+^CD8^+^ T cells producing INF-γ and/or TNF-α. I-J. Bar diagrams summarize the count of CD45.2^+^CD8^+^ cells producing IFN-γ and TNF-α (I) or IFN-γ (J). Data represents Mean ± SEM; ***p<0.001; **p<0.005; *p<0.05 and ns (p>0.05)- not significant (One-way ANOVA).

**Figure S5.**
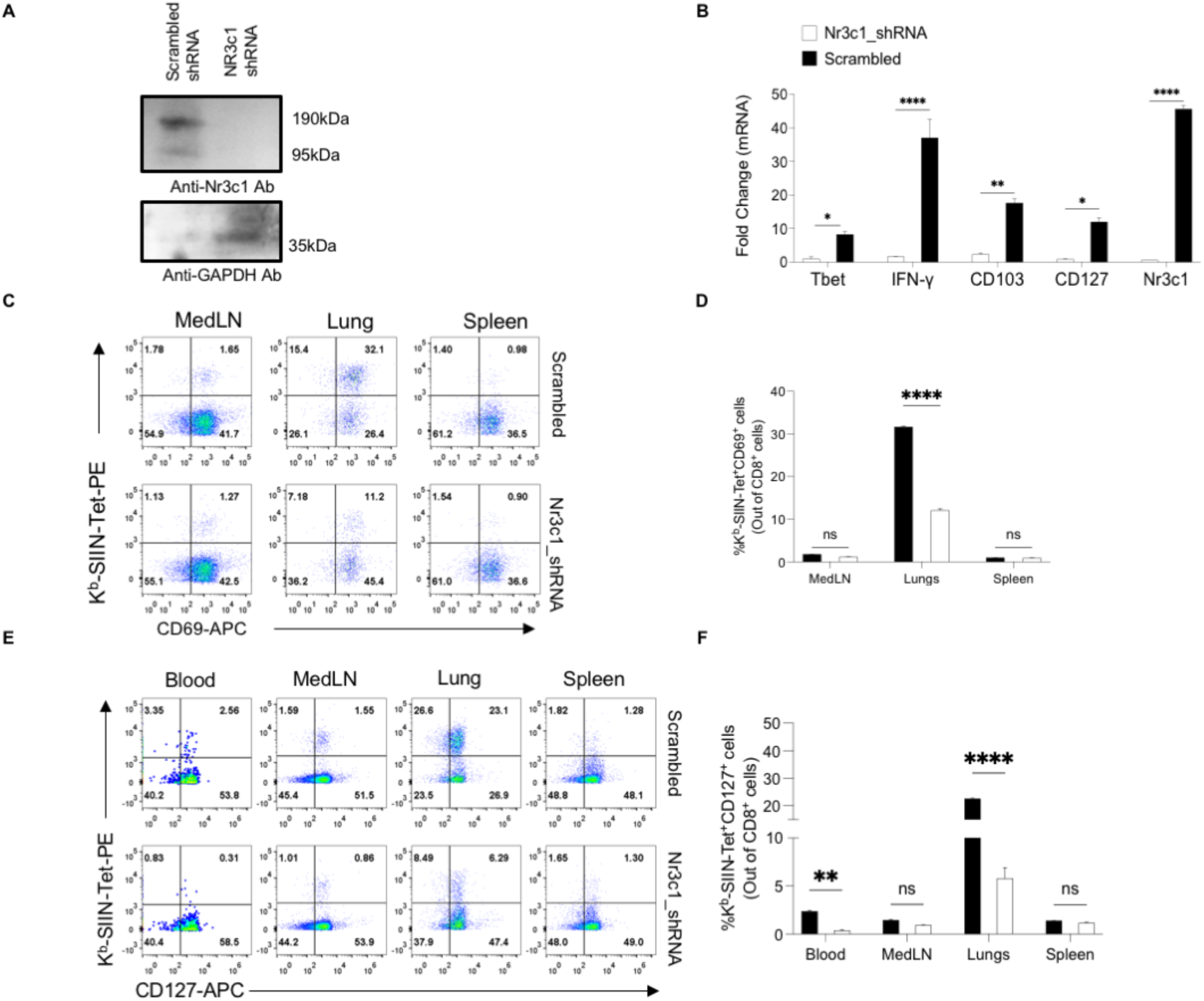
Phenotype of cells lacking the glucocorticoid receptor. Enriched OT1 cells were *in vitro* stimulated and transduced with retroviruses coding the shRNA against *nr3c1* or the scrambled sequence. 5 days post transduction, the cells were used for RNA isolation and cell lysate preparation. Subsequent analysis was performed to analyse the expression of various molecules by qRT-PCR and a western blot was performed to confirm knockdown of *nr3c1*. A. The immunoblot image to confirm *nr3c1* knockdown. B. Bar diagrams summarizing the comparison of the relative expression of the molecules mentioned by qRT-PCR are shown. C-F. *In vivo* phenotypic analysis of cells lacking *nr3c1*. C. Representative FACS plots show the proportion of CD69^+^Tetramer^+^ cells in different compartments. D. Bar diagram summarizing the frequency of CD69^+^Tetramer^+^ cells in the two groups. E. Representative FACS plots for the analysis of CD127^+^Tetramer^+^ cells in the compartments mentioned are shown. F. Bar diagram summarizing the frequency of CD127^+^Tetramer^+^ cells in the compartments mentioned is shown. Data represents Mean ± SEM; ***p<0.001; **p<0.005; *p<0.05 and ns (p>0.05)- not significant (Two-way ANOVA).

**Figure S6.**
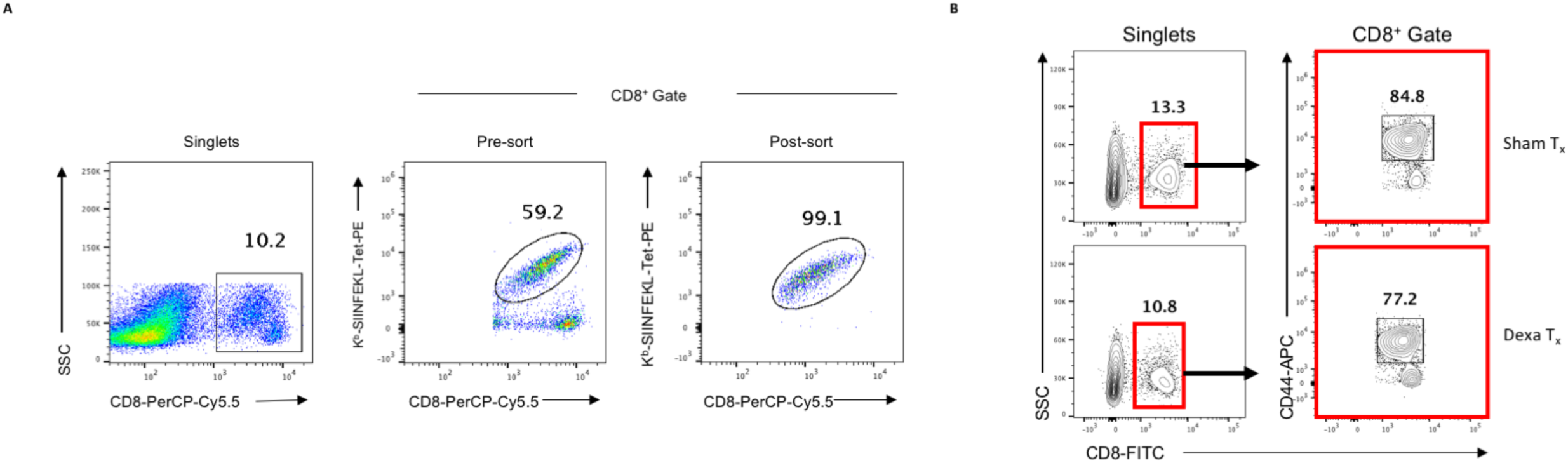
Sorting strategy for Tetramer positive (A) and CD8^+^CD44^+^ cell sorting (B).

**Figure S7.**
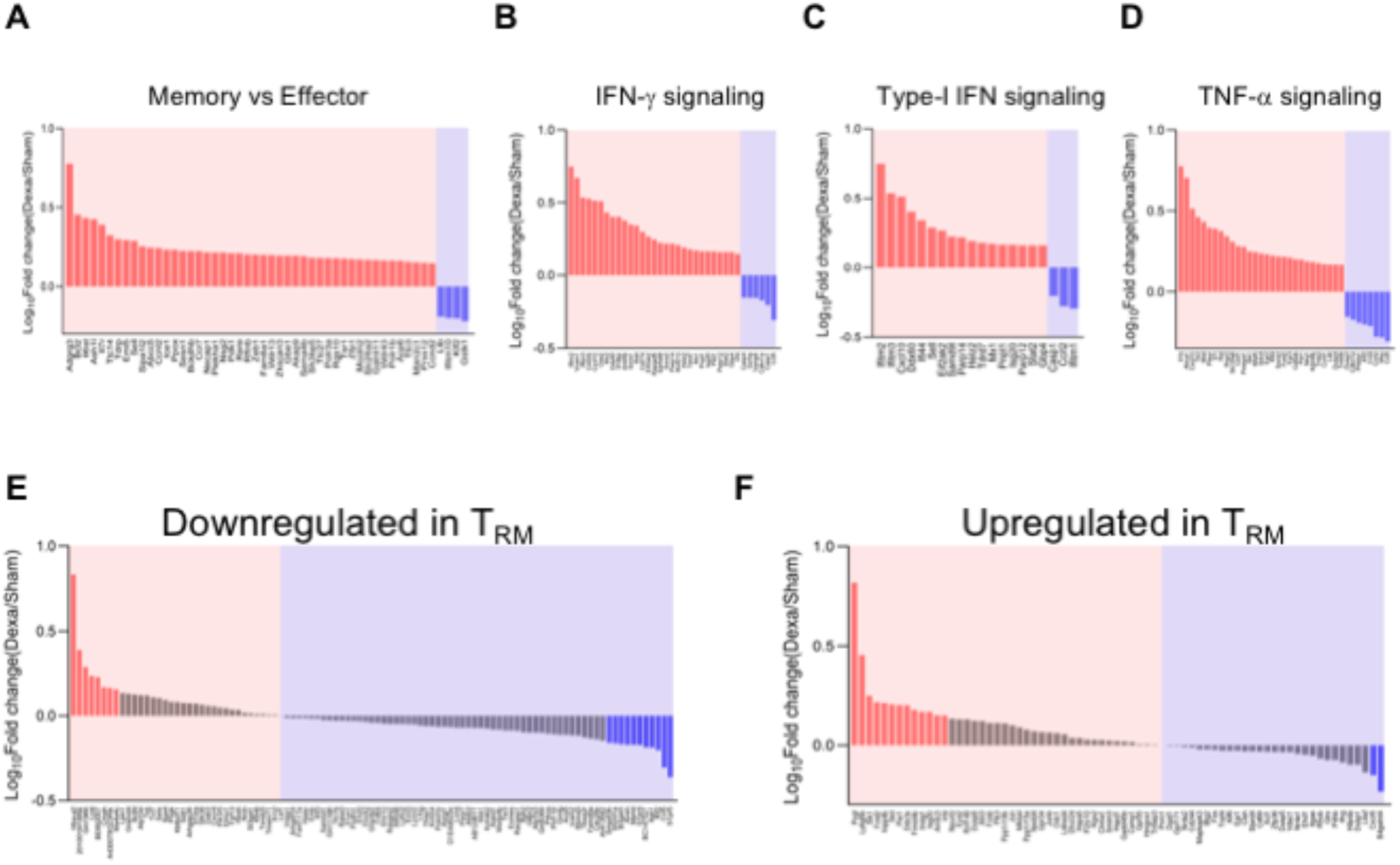
Dexamethasone treated CD8^+^ T cells exhibit a transcriptional program skewed towards tissue resident memory CD8^+^ T cells. Gene set enrichment analysis was performed with gene sets obtained from immunesigdb of the molecular signatures database. A-F. Log_10_Fold change (Dexa T_x_/ Sham T_x_) for genes known to be upregulated in memory cells as compared to effector cells (A), genes upregulated in response to IFN-γ (B), IFN-α (C) and TNF-α (D) signaling and genes known to be downregulated (E) and upregulated (F) in T_RM_ are plotted.

**Figure S8.**
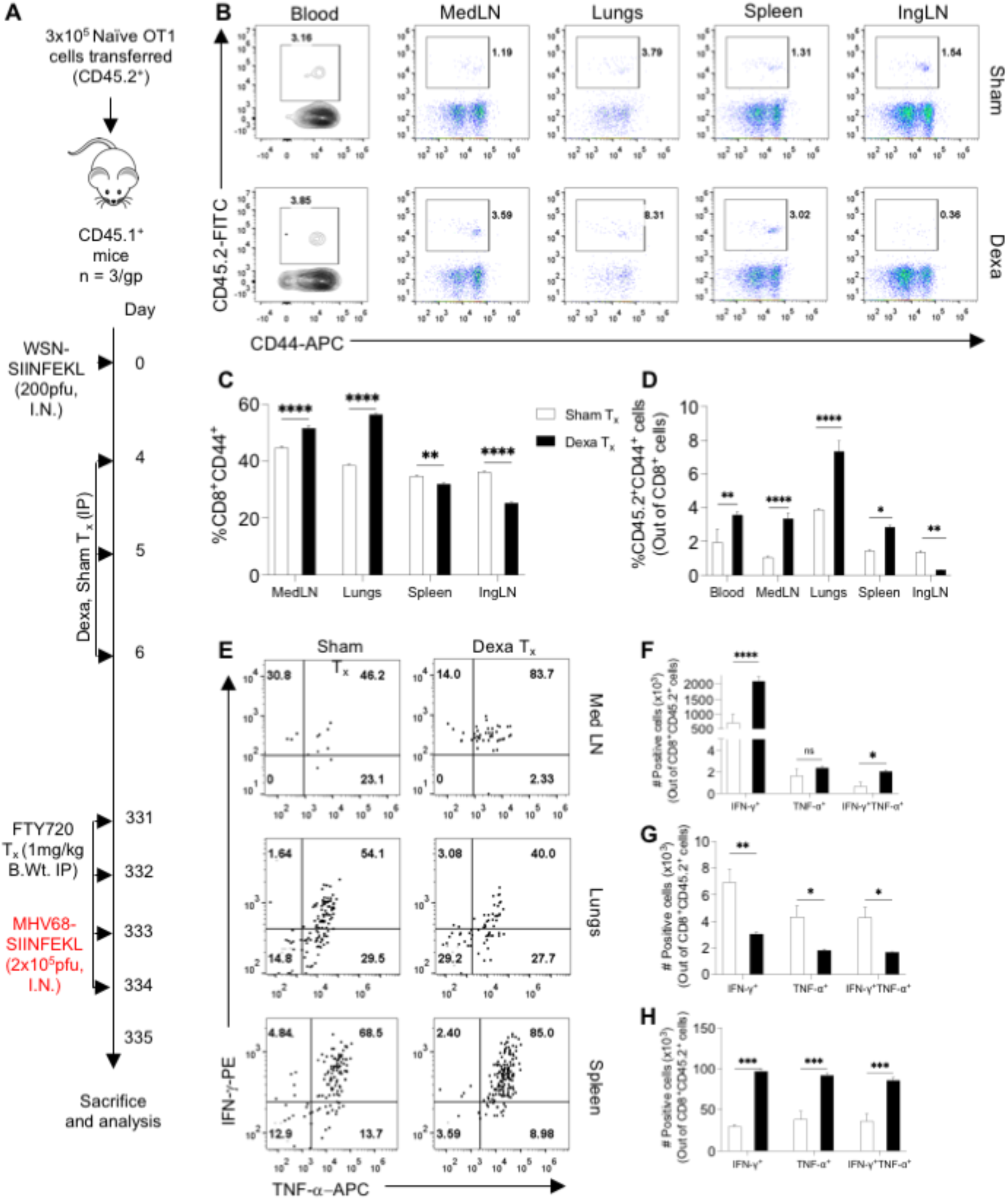
A transient corticosteroid therapy enhances local immunity. A. A schematic of the experiments is shown. Enriched OT1 cells were adoptively transferred into naïve sex matched congenic CD45.1^+^ mice which were than infected intranasally with 200pfu of WSN-SIINFEKL and treated with dexamethasone or sham at 4,5 and 6 dpi. 1mg/kg body weight FTY720 treatment was initiated from 331 dpi and the animals were given a heterologous challenge with MHV68-SIINFEKL at 334dpi. The animals were sacrificed 2 days post recall and subjected to cellular analysis. B. Representative FACS plots for the frequency of Donor cells in the different compartments are shown. C, D. Frequencies of activated (CD8^+^CD44^+^, C) and donor (CD45.2^+^, D) cells in the different compartments are summarized by bar diagrams. The functionality of the OT1 cells upon recall were analyzed by ICCS assays. E. Representative FACS plots show the frequencies of IFN-γ and TNF-α producing OT1 cells following *in vitro* peptide pulse. F-H. Bar diagrams summarize the absolute counts of single (IFN-γ or TNF-α) and double (IFN-γ and TNF- α) cytokine producing cells in the draining medLN, F), lungs (G) and spleen (H). Data represents Mean ± SEM; ***p<0.001; **p<0.005; *p<0.05 and ns (p>0.05)- not significant (Two-way ANOVA).

**Supplementary table 1.**
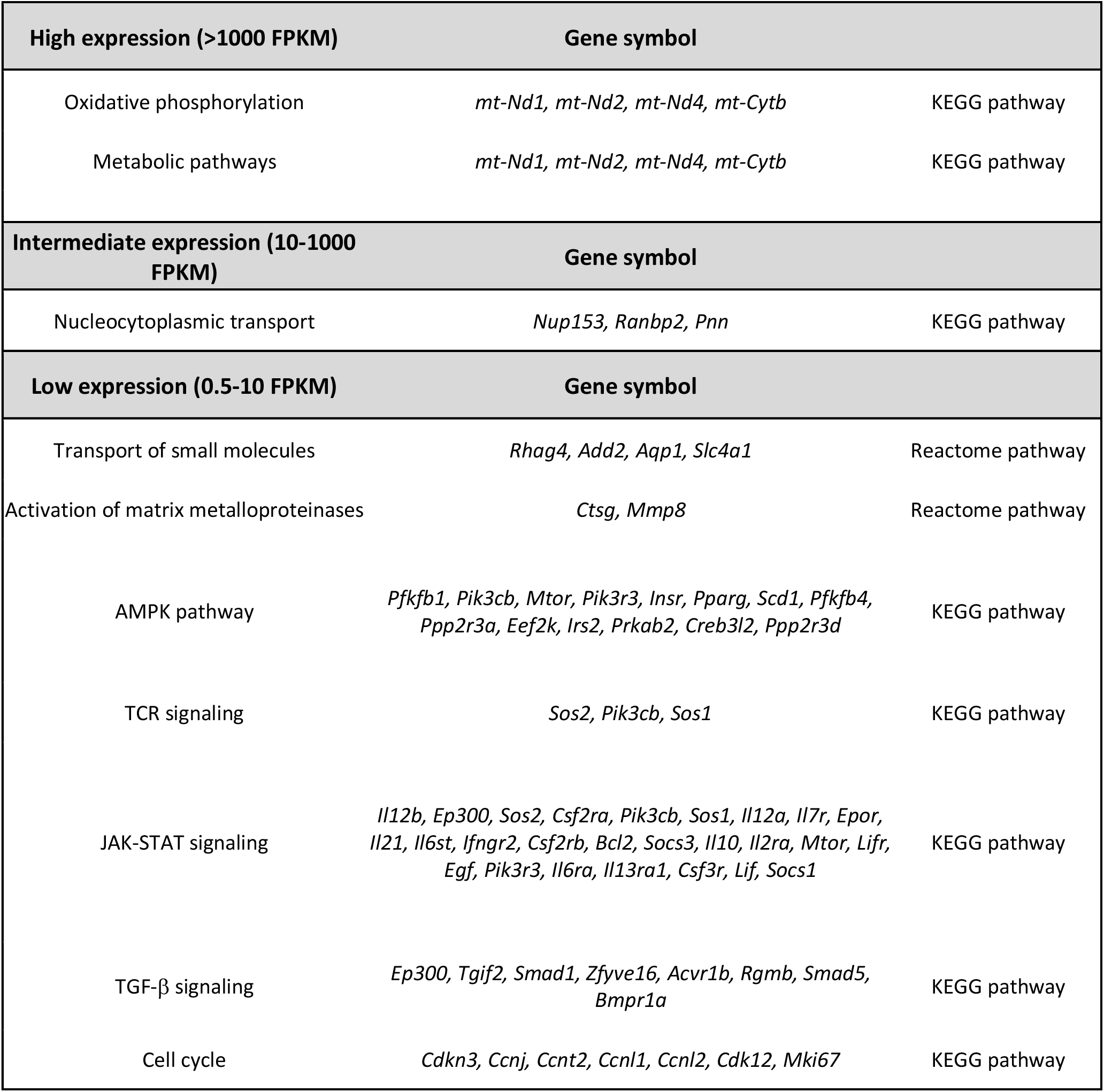
Pathway enrichment analysis

